# Structural basis for selective stalling of human ribosome nascent chain complexes by a drug-like molecule

**DOI:** 10.1101/315325

**Authors:** Wenfei Li, Fred R. Ward, Kim F. McClure, Stacey Tsai-Lan Chang, Elizabeth Montabana, Spiros Liras, Robert Dullea, Jamie H. D. Cate

## Abstract

Small molecules that target the ribosome generally have a global impact on protein synthesis. However, the drug-like molecule PF-06446846 (PF846) binds the human ribosome and selectively blocks the translation of a small subset of proteins by an unknown mechanism. In high-resolution cryo-electron microscopy (cryo-EM) structures of human ribosome nascent chain complexes stalled by PF846, PF846 binds in the ribosome exit tunnel in a newly-identified and eukaryotic-specific pocket formed by the 28S ribosomal RNA (rRNA), and redirects the path of the nascent polypeptide chain. PF846 arrests the translating ribosome in the rotated state that precedes mRNA and tRNA translocation, with peptidyl-tRNA occupying a mixture of A/A and hybrid A/P sites, in which the tRNA 3’-CCA end is improperly docked in the peptidyl transferase center. Using mRNA libraries, selections of PF846-dependent translation elongation stalling sequences reveal sequence preferences near the peptidyl transferase center, and uncover a newly-identified mechanism by which PF846 selectively blocks translation termination. These results illuminate how a small molecule selectively stalls the translation of the human ribosome, and provides a structural foundation for developing small molecules that inhibit the production of proteins of therapeutic interest.

## Main Text

Most compounds that target the ribosome affect a large number of mRNAs through general inhibition of translation initiation or elongation ^1-3^. Furthermore, these compounds almost exclusively act as broad-spectrum inhibitors, displaying little sequence specificity ^4^. Many studies have revealed how these compounds bind the translating ribosome and inhibit its function ^5-9^. Recently, we described the small molecule PF846 that selectively blocks the translation of individual mRNAs by the human ribosome ^10^. However, a mechanism to account for the ability of PF846 to selectively inhibit human protein synthesis remains unknown.

PF846 blocks production of proprotein convertase subtilisin/kexin type 9 (PCSK9)–an important target for regulating plasma low-density lipoprotein cholesterol levels–by interfering with the elongation phase of translation ^10,11^. PF846 selectively stalls the ribosome on very few translated protein nascent chains, generally early in their formation and with no clear sequence pattern ^10^. To gain insight into the mode of action of PF846 in targeting the human ribosome and specific nascent chains, and to identify principles for future drug development, we used single-particle cryo-EM to determine structures of PF846 stalled ribosome nascent chain (RNC) complexes. We also employed randomized mRNA libraries to identify sequence preferences for PF846-mediated translation stalling.

Since the stalling sequences in the few proteins affected by PF846 mostly occur near the N-terminus and would therefore reside in the ribosome exit tunnel ^10^, we first identified conditions for affinity purification of stable PF846-stalled RNCs from *in vitro* translation reactions in human cell extracts. The calcium-dependent cell-cell adhesion glycoprotein Cadherin-1 (CDH1) ^12^ is stalled near its C-terminus by PF846, enabling the extension of the peptide N-terminus beyond the confines of the ribosome exit tunnel ^13^. Appending an N-terminal affinity tag followed by the CDH1-V domain and the nascent chain sequences targeted by PF846 did not disrupt the ability of PF846 to stall translation and allowed formation and purification of stable PF846-stalled RNCs (**Fig. 1a, b, Extended Data Fig. 1**).

**Fig 1.**
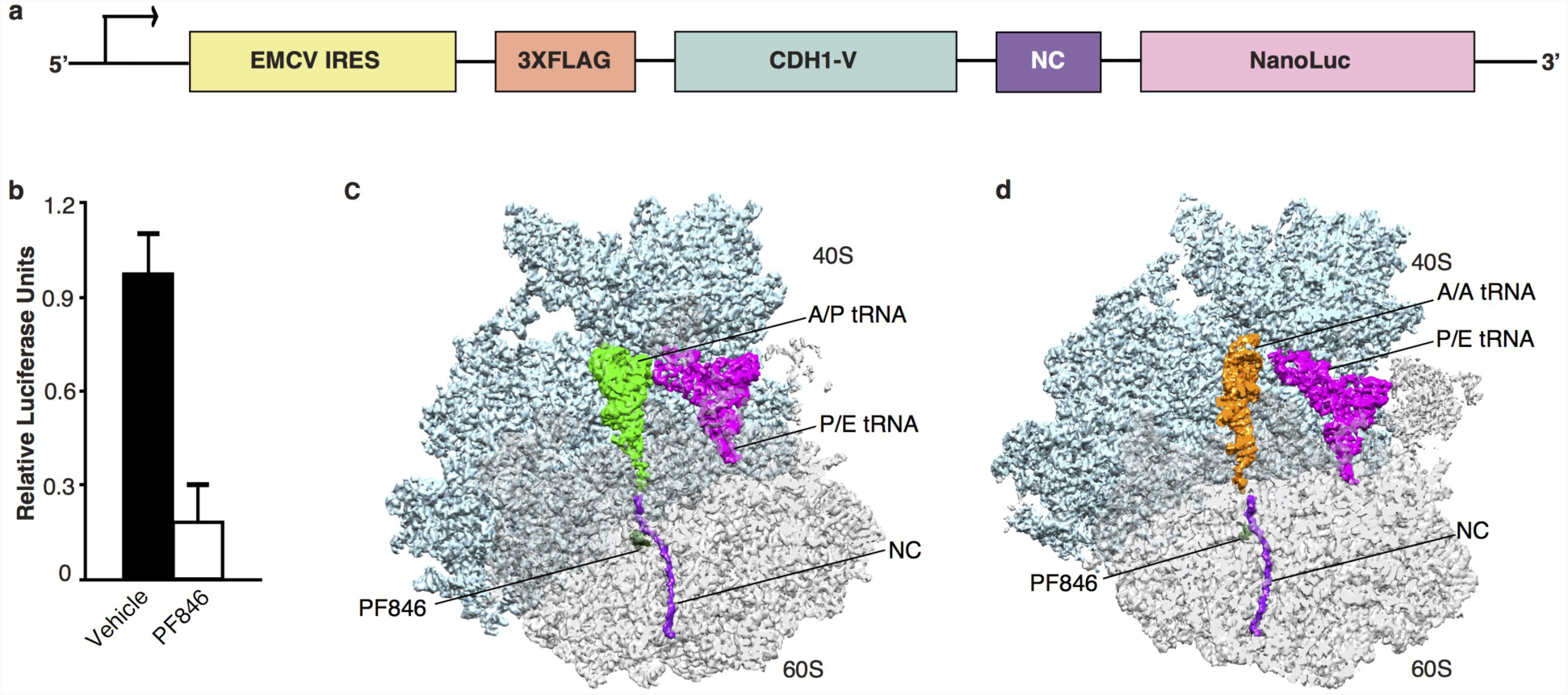
Structural analysis of PF846 stalled RNCs. **a**, Schematic representation of the DNA construct used to prepare PF846 stalled RNCs. **b**, Stalling of the chimeric PCSK9 nascent chain in *in vitro* translation reactions in the absence (black bar) or presence (white bar) of PF846. **c**, Cryo-EM structure of the stalled CDH1-RNC in the rotated state showing the A/P tRNA (green), P/E tRNA (magenta), CDH1 nascent chain (purple) and PF846 (dark green) with small and large subunit colored in cyan and grey, respectively. **d**, Structure of the stalled CDH1-RNC in the rotated state with A/A tRNA (orange), and P/E tRNA (magenta).

### Overall structures of PF846 stalled RNCs

To investigate how PF846 stalls specific nascent chain sequences in the human ribosome, we used cryo-EM to determine structures of PF846-stalled CDH1, PCSK9 and USO1 ribosome nascent chain complexes (CDH1-RNC, PCSK9-RNC and USO1-RNC, respectively) (**Extended Data Figs. 2, 3, and 4, Extended Data Tables 1, 2, 3, and 4**). In both the CDH1-RNC and PCSK9-RNC samples, particle sorting of the purified RNCs revealed a major population of ribosomes in the rotated state that precedes mRNA and tRNA translocation ^14^, containing predominantly hybrid A/P site peptidyl-tRNA (i.e. nascent chain-tRNA, or NC-tRNA), but with some rotated-state RNCs containing A/A site NC-tRNA. The USO1-RNC also adopts the rotated state, although the positioning of the A-site tRNA in the A/P or A/A site was less well defined. All three RNCs also included tRNA in the hybrid P/E site and PF846 bound in the ribosome exit tunnel (**Fig. 1c, d, Extended Data Fig. 5**). The following analyses describe the CDH1-RNCs due to the higher-quality of the CDH1-RNC cryo-EM maps, although we observe similar behavior for the PCSK9-RNC and USO1-RNC samples, as shown in the Extended Data Figures.

**Fig 2.**
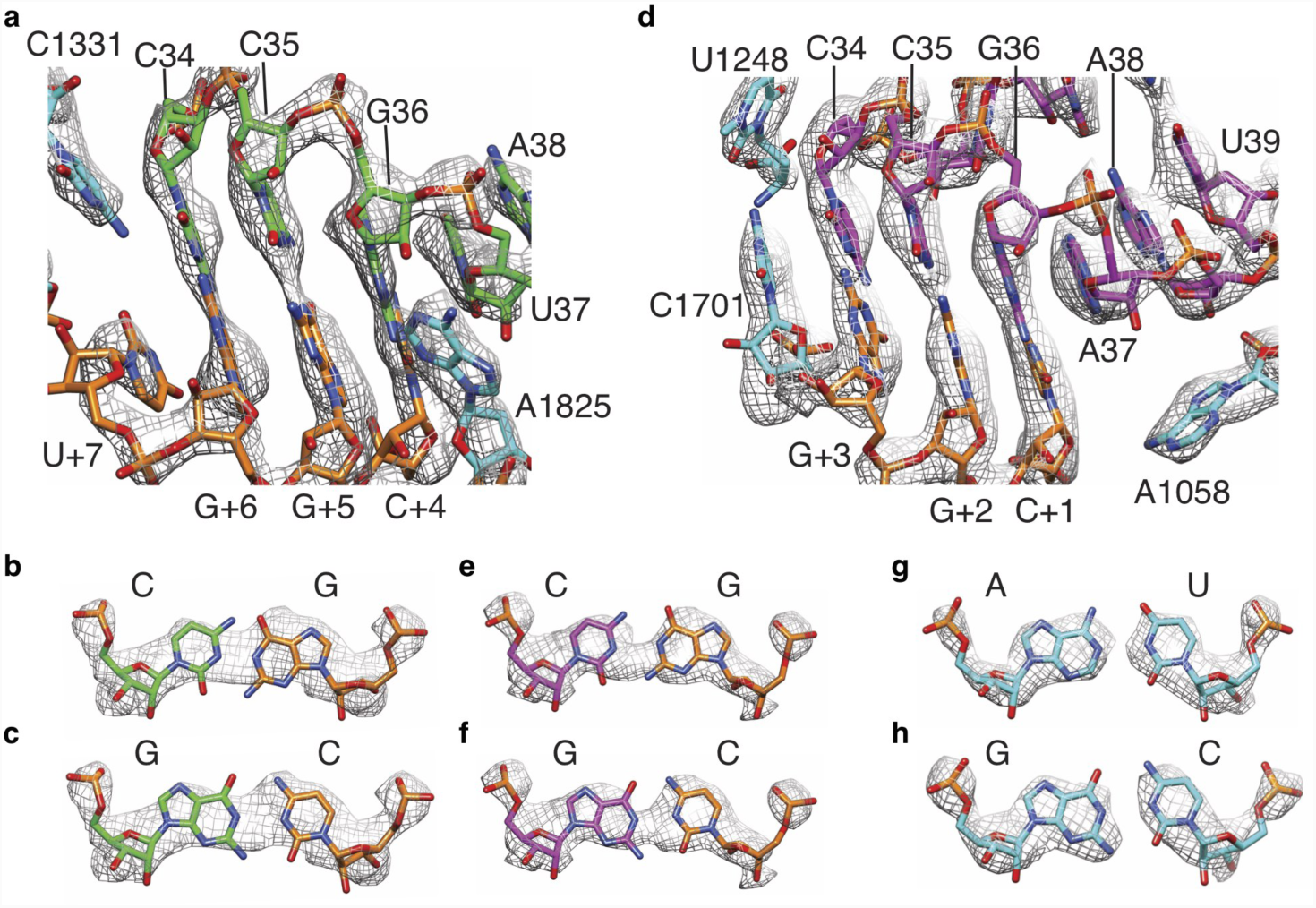
Analysis of the interactions of A/P tRNA and P/E tRNA with mRNA in the PF846 stalled CDH1-RNC. **a**, Density for A-site mRNA (orange) and anticodon stem-loop (ASL) region of A/P tRNA (green). **b-c**, Models for a G-C or C-G base pair between A/P tRNA anticodon nucleotide 34 and mRNA codon nucleotide +6 (3rd nucleotide of codon in the A site) in the observed density. **d**, Density for P-site mRNA (orange) and ASL of P/E tRNA (magenta). **e-f**, Models for a G-C or C-G base pair between P/E tRNA anticodon nucleotide 34 and mRNA codon nucleotide +3 (3rd nucleotide of codon in the P site) in the observed density. **g-h**, Observed density for representative A-U (A1052 and U1066) and G-C (G1065 and C1053) base pairs in 18S rRNA (cyan) in the 40S head and platform domains, near the P site.

**Fig 3.**
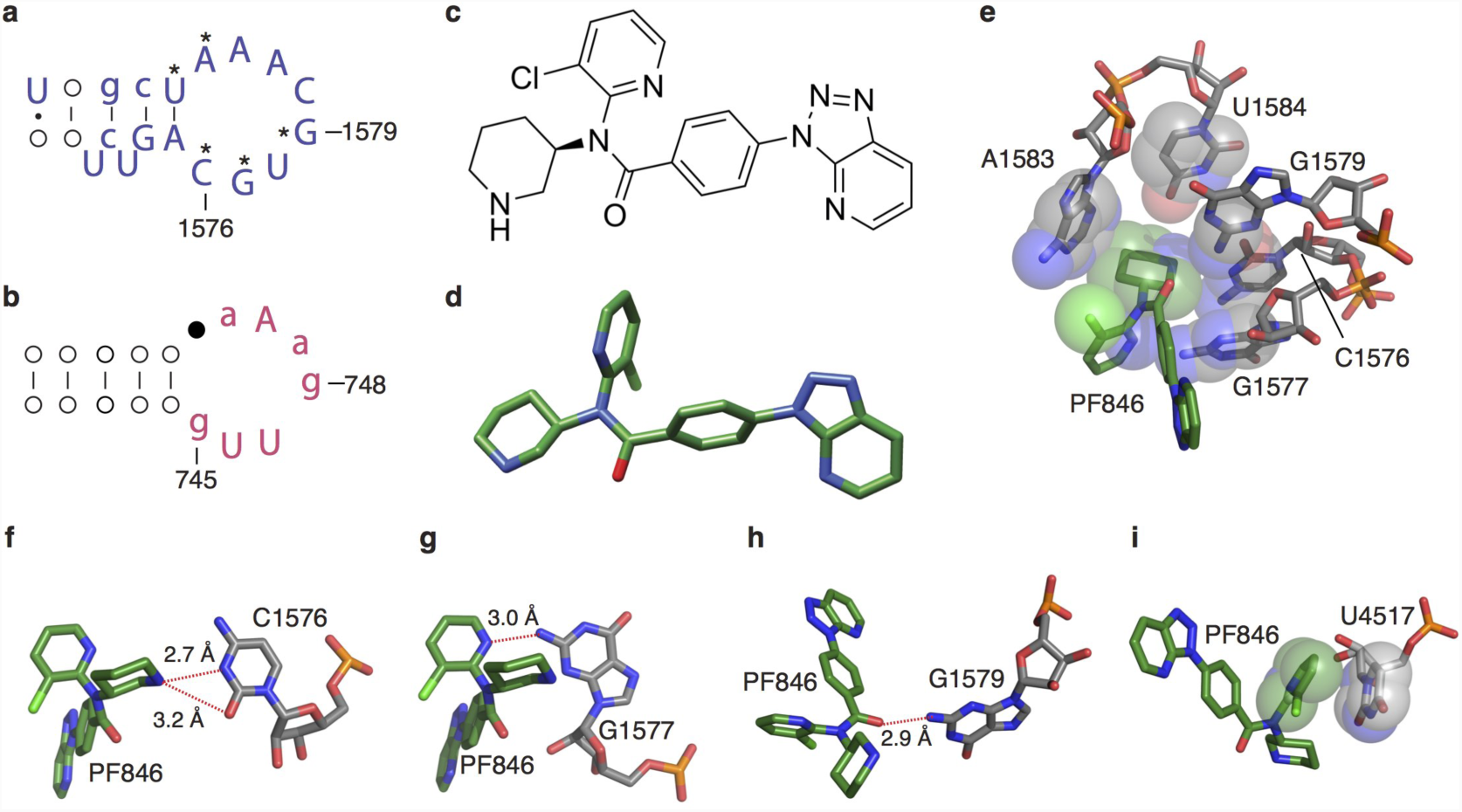
Interactions of PF846 with the human ribosome. **a-b**, Phylogenetic analysis of the 28S rRNA binding pocket in eukaryotes (**a**) and bacteria (**b**). Nucleotides that interact with PF846 are labeled with an asterisk (*). Nucleotides with capital and small letters (AUCG, aucg), are over 98% conserved or 90%-98% conserved, respectively. **c-d**, PF846 chemical and three-dimensional structure, respectively. **e**, 28S rRNA residues with direct interactions with the piperidine ring of PF846. Surfaces represent the van der Waals radii of the C, N and O atoms. **f-h**, Interactions of PF846 and nucleotides from 28S rRNA. **i**, Interaction of the chloro-pyridine ring with U4517, with van der Waals surfaces as in **e**.

**Fig 4.**
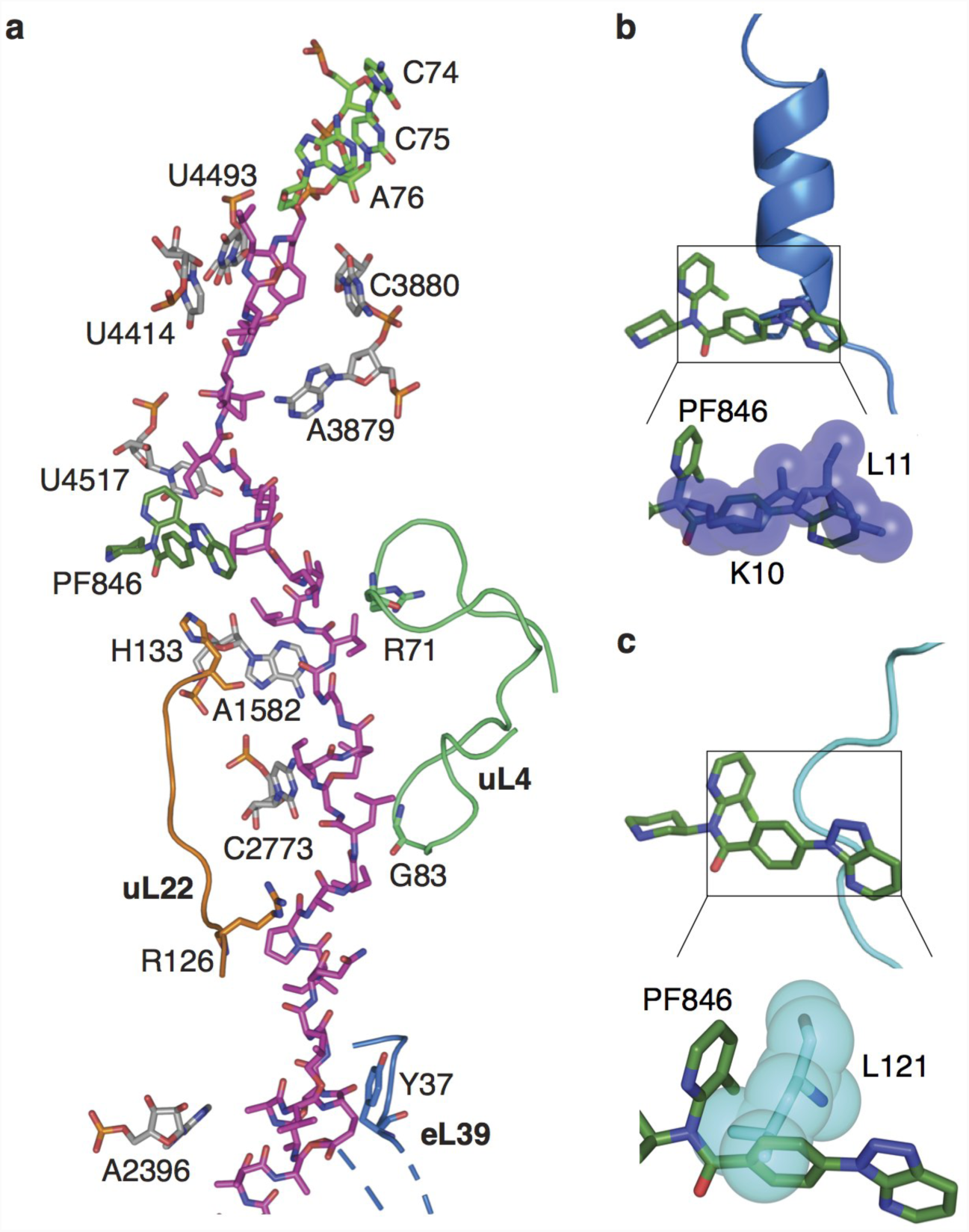
CDH1 nascent chains and comparisons with other stalled peptides in the ribosome exit tunnel. **a**, Molecular model of CDH1 nascent chain (magenta) following the 3’-CCA end of A/P tRNA (green) in the ribosome exit tunnel with 28S rRNA (grey), uL22 (orange), uL4 (light green) and PF846 (dark green) highlighted. **b-c**, Superposition of hCMV ^18^ (blue) and SRP ^19^ (cyan) stalled nascent chains within the exit tunnel, showing the predicted steric clashes with PF846 (dark green), using van der Waals surfaces.

The CDH1-RNC complexes with hybrid A/P site NC-tRNA or with A/A site NC-tRNA yielded cryo-EM maps with an average resolution of 3.0 Å and 3.9 Å, respectively (**Fig. 1c, d, Extended Data Fig. 2**). The hybrid A/P and P/E NC-tRNAs each represent a mixture of tRNAs, as inferred from the EM map density for nucleotide bases in the codon-anticodon base pairs (**Fig. 2, Extended Data Fig. 6**), consistent with the clustering of stall sites spanning multiple codons observed by ribosome profiling ^10^. In the CDH1-RNC and PCSK9-RNC samples, we also observed a small population of PF846-stalled RNCs in the non-rotated state bearing a P/P-site NC-tRNA that exists after peptide bond formation (**Extended Data Figs. 2, 3, 5**). Interestingly, this population appeared in higher abundance when the PCSK9-RNC sample was purified using a short isolation time prior to making cryo-EM grids (**Extended Data Fig. 3d-e**). These results suggest that the non-rotated RNC is a transient state during PF846 mediated translation stalling. Collectively, our findings suggest that PF846 preferentially stalls RNCs in the rotated state, while the non-rotated state with P/P-site NC-tRNA can be resolved during translation in the presence of PF846.

**Fig 5.**
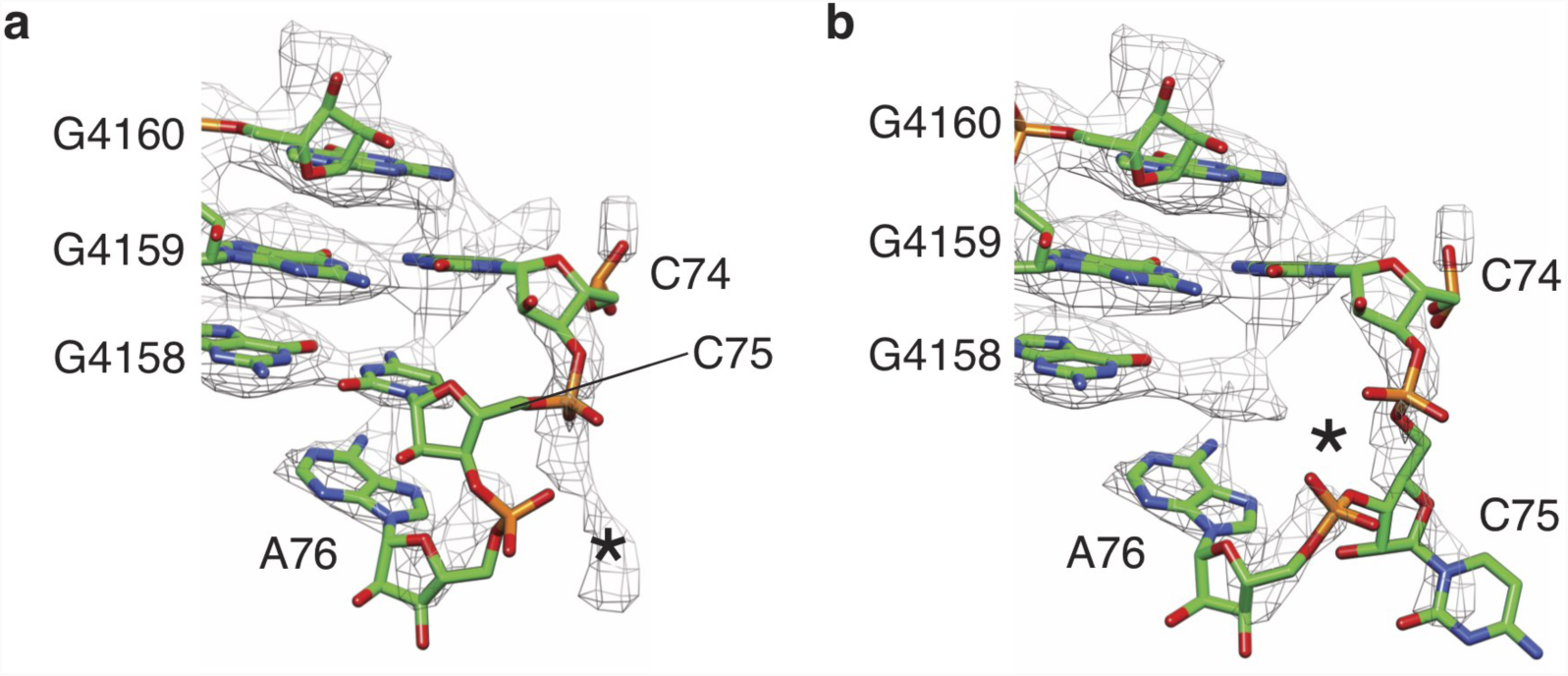
Disrupted based pairing between tRNA 3’-CCA end and 28S rRNA in the ribosomal large subunit P site. **a**, Poor positioning of A/P NC-tRNA 3’-CCA end near the ribosomal P-loop. High-resolution structure of the 3’-CCA end of peptidyl-tRNA (PDB: 1vy4) ^35^ is docked as a rigid body, using nucleotides from 28S rRNA in the PTC as reference. **b**, Nucleotide C75 in A/P NC-tRNA modeled as flipped to nearby density without pairing with the P-loop (starting model from PDB: 1vy4).

### Eukaryotic-specific small molecule binding pocket

In the stalled RNCs, PF846 occupies a binding pocket in a newly-identified groove of the ribosomal exit tunnel formed by universally conserved 28S rRNA nucleotides ^15^ (**Fig. 3**) and a stretch of the stalled nascent chain, best seen in the rotated-state RNCs with A/P NC-tRNAs. PF846 was fit as a rigid body to the density by evaluating its X-ray structures ^10^ and low energy conformations based on quantum mechanical assessment of accessible torsion angles (**Extended Data Fig. 7, Extended Data Table 5**), followed by adjustments of the chloropyridine and aryl rings. Consistent with the ability of PF846 to bind to non-translating 80S ribosomes ^10^, contacts with 28S rRNA make up the bulk of the small molecule interactions with the ribosome (**Fig. 3**). The piperidine ring extends into the rRNA binding pocket, contacting nucleotides C1576, G1577, G1579, A1583 and U1584 (**Fig. 3e, f**). The chloropyridine and amide linkage also form hydrogen bonds with highly conserved nucleotides G1577 and G1579 (**Fig. 3g, h**). Additionally, the interaction is stabilized by U4517, which forms a stacking interaction with the chloro-pyridine ring (**Fig. 3i**). To explore the structural basis for PF846 action on eukaryotic but not bacterial ribosomes ^10^, we aligned the small molecule-binding pocket with the corresponding region in the *E. coli* ribosome (**Fig. 3a, b, Extended Data Fig. 8**). Phylogenetic analysis revealed key changes to the nucleotide pattern in 23S rRNA in the large ribosomal subunit of bacteria ^15^ (**Fig. 3a, b**). Whereas nucleotides C1576, G1577 and G1579 in the human ribosome are highly complementary to the conformation of PF846 (**Fig. 3e-h**), nucleotides *N1*-methyl-G745 and ψ746 in bacterial 23S rRNA would clash with the compound (**Extended Data Fig. 8c, d**). Furthermore, whereas U4517 in the human ribosome stacks on the chloropyridine ring in PF846 (**Fig. 3i**), the corresponding nucleotide in bacterial 23S rRNA (U2609) would sterically clash with PF846 (**Extended Data Fig. 8e**). Taken together, these distinctions in nucleotide identity and positions help explain PF846’s specificity for eukaryotic ribosomes.

### Analysis of stalled nascent chains within the exit tunnel

The triazolopyridine ring system of PF846 (**Fig. 3c, d**) faces the ribosome tunnel and is the only part of the molecule with direct interaction with the stalled nascent chain (**Fig. 4**, **Extended Data Figs. 9, 10**). Notably, the triazolopyridine moiety has incomplete density (**Extended Data Fig. 7**) that may be caused by the flexibility of this region during nascent chain elongation. Interestingly, the density for the nascent chains in the RNCs with A/P NC-tRNA was mostly well defined in the ribosome exit tunnel with a local resolution of ∼4-5 Å, although they are likely comprised of different sequences superimposed on each other corresponding to the cluster of stalled states spanning multiple mRNA codons (**Fig. 2, Extended Data Fig. 6**) ^10^. All three of the nascent chains (CDH1, PCSK9, USO1) have similar geometry and adopt a predominantly extended conformation in the exit tunnel ^16^ (**Extended Data Figs. 9, 10**). The nascent chain spans ∼9 residues between the C-terminus of the nascent chains bonded to the 3’-CCA end of A/P site tRNA in the peptidyl transferase center (PTC) and the small molecule binding site (**Fig. 4a, Extended Data Fig. 10**). These residues engage in multiple interactions with nucleotides in 28S rRNA of the 60S subunit, including U4493, U4414, and A3879 (**Fig. 4a**, **Extended Data Fig. 10**). Past the compound-binding pocket (i.e. in the N-terminal direction), the nascent chain contacts ribosomal proteins uL4 and uL22, along with A1582 and C2773 in 28S rRNA (**Fig. 4a, Extended Data Fig. 10**). These contacts distal from the peptidyl transferase center are likely important for the stalling induced by PF846, based on N-terminal deletion analysis of the PCSK9 sequence ^10^.

Most reported RNC structures have “kinks” in the nascent chain residing in the ribosome exit tunnel proximal to the PTC ^17^. The CDH1, PCSK9 and USO1 nascent chains also have a kink in their structures between the PTC active site and PF846-binding pocket (**Fig. 4a, Extended Data Figs. 9, 10**), but these adopt a different geometry when compared to stalled human cytomegalovirus (hCMV) and signal recognition particle (SRP) nascent chains ^18,19^ (**Fig. 4b, c**). In the PF846-stalled RNC structures, the “kinks” assume more acute angles, to make enough space for the bound small molecule (**Fig. 4a, Extended Data Figs. 9, 10**), whereas the hCMV and SRP NC positions would occlude PF846 binding (**Fig. 4b, c**).

### Destabilized interaction between the A/P-tRNA and the P-loop in PF846 stalled RNCs

PF846 stalls translation of the ribosome predominantly in the rotated state that precedes mRNA and tRNA translocation to the next codon, presumably impeding the action of eEF2, the GTPase which promotes the translocation reaction ^20^. In bacterial translation, which is better understood ^21,22^, translocation proceeds through a series of ribosome rotated states, in which the peptidyl-tRNA body and 3’-CCA end move independently from one another, transiting from an A/A site with 3’-CCA end base pairing with the large subunit rRNA A-loop, to an A/A site in the rotated state in which the 3’-CCA end base pairing with the large subunit rRNA A-loop is retained, and finally to an A/P site with 3’-CCA base pairing with the P-loop ^23^. At this point, EF-G (the bacterial homologue of eEF2) can catalyze the remaining conformational changes required for complete translocation of the mRNA and tRNAs ^21,22^. In the PF846-stalled complexes, we observe pre-translocation RNC populations with A/A and A/P NC-tRNAs. However, the A/P-tRNA exhibits flexible base pairing of the 3’-CCA end with the ribosomal P-loop (**Fig. 5a, b, Extended Data Fig. 11**). We propose that PF846 inhibits rapid translocation of specific nascent chains by eEF2, by perturbing the path of the NC remotely from the PTC and thereby preventing the peptidyl-tRNA from binding stably in the hybrid A/P site, which requires the tRNA 3’-CCA end to properly base pair with 28S rRNA nucleotides in the P-loop.

### Sequence Determinants of PF846-induced stalling

The few NC sequences stalled by PF846 as previously identified by ribosome profiling do not reveal a simple amino acid “motif” responsible for stalling ^10^. We therefore used *in vitro* translation reactions to identify the sequence determinants in the NC required for PF846-induced stalling of RNCs. To assay the sequence-dependence along the NC, we first used a poly-asparagine scan, in which we replaced 3-4 amino acids in the stalling sequence at a time. We used asparagine scanning as opposed to alanine scanning used previously ^10^ to maintain a rough approximation to the size distribution of the amino acids in the NC, while introducing a hydrophilic side chain. In both the CDH1 and PCSK9 stalling sequences, replacement of leucine- and isoleucine-rich stretches of hydrophobic amino acids predicted to be in the PTC-proximal region of the ribosome exit tunnel abrogated PF846-induced stalling (**Extended Data Fig. 12**). By comparing to the ribosome footprinting ^10^ and structural analyses (**Fig. 4**, **Extended Data Figs. 9, 10**), the most important sequences for CDH1 span the NC from the PTC to just beyond the PF846 binding site. For PCSK9, the hydrophobic stretch of amino acids identified to be important for PF846-induced stalling is likely to be positioned just beyond the PF846 binding site in the exit tunnel (**Extended Data Fig. 10b**).

The Asn-scanning experiments above revealed NC sequences sensitive to mutation, but did not reveal the spectrum of amino acid sequences that can be stalled by PF846. To identify the range of amino acid sequences that can be stalled by PF846 in an unbiased manner, we used an mRNA library-based approach followed by deep sequencing, analogous to ribosome display ^24^. We randomized four NC amino acids at a time near the PTC and near the PF846 binding site, by introducing (NNK)_4_ sequences (N, any nucleotide; K, G or U) into mRNAs encoding the CDH1 stalling sequence (**Fig. 1a**, **Extended Data Fig. 13a**). We used these mRNA libraries in *in vitro* translation reactions, in the presence or absence of PF846, and then used the 3X-FLAG tag at the N-terminus of the NC to pull down stalled RNCs, which would retain the mRNAs encoding the stalled NC sequences. Next, we used RNaseH to eliminate mRNAs translated by elongating ribosomes, i.e. ribosomes in the middle of the luciferase ORF downstream of the stall sequence (**Extended Data Fig. 13b**). The mRNAs isolated in this manner were then converted into a DNA library and analyzed by deep sequencing.

In the mRNA library with randomized sequences near the PTC, we observed strong enrichment for PF846-sensitive stalling sequences strikingly different from the original CDH1 sequence (**Fig. 6**, **Extended Data Table 6**). In particular NC positions –2 and –4 from the predicted stall site are enriched in histidine rather than leucine (**Fig. 6a**, inset). Other polar and charged amino acids are also enriched in the PF846-sensitive NCs. We validated two of the newly-identified stalling sequences using *in vitro* luciferase assays. We found PF846 stalled HYHS and RSCK sequences with a similar IC50 compared to LLLL in the original CDH1 sequence (**Fig. 6b**, **Extended Data Fig. 13c**), with the HYHS sequence being more potently stalled at high PF846 concentrations (**Fig. 6b**). Both sequences were also enriched in stalled RNCs, as determined by Western blotting (**Fig. 6d, Extended Data Fig. 13d**). Surprisingly, we observed no enrichment of PF846-stalling dependent sequences in the mRNA library encoding randomized NCs in the ribosome exit tunnel near the PF846 binding site (**Extended Data Fig. 13e**). This is in striking contrast to the Asn-scanning results, in which mutation of NC sequences near the PF846 binding site disrupted stalling. This is likely due to the fact that Asn-scanning introduced negative determinants of PF846-dependent stalling, whereas enrichment from the mRNA libraries revealed positive determinants of stalling.

**Fig 6.**
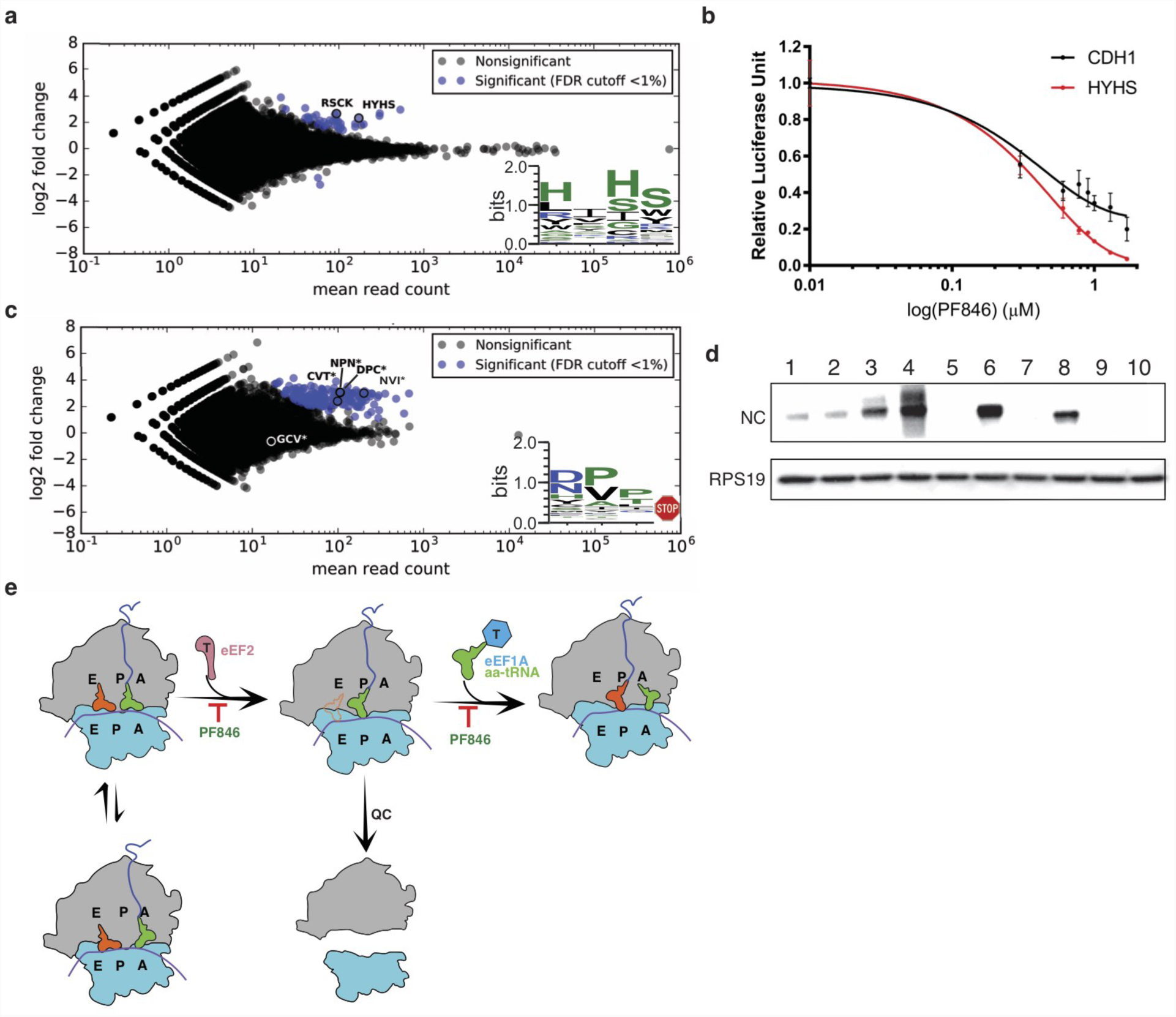
Selections of PF846-dependent stalling sequences from randomized mRNA libraries. **a**, MA plot of sequences enriched for PF846-induced translation elongation stalling, compared to translation reactions in the absence of PF846. Sequences were enriched from the mRNA library of CDH1-derived nascent chains with four amino acid positions randomized near the predicted stall site in the PTC. Log_2_-fold enrichment of sequences is plotted against total read count, for experiments carried out in duplicate. Blue, sequences enriched with an adjusted *P*-value < 0.01. Inset, consensus sequence logo for enriched sequences. **b**, Luciferase activity of translation reactions with luciferase reporters containing the WT CDH1 stalling sequence or with the HYHS motif near the PTC, as a function of PF846 concentration. Reactions were carried out in triplicate, with standard deviation shown. **c**, MA plot of sequences enriched for PF846-induced inhibition of translation termination, compared to translation reactions in the absence of PF846. Sequences were enriched from the mRNA library of CDH1-derived nascent chains with four amino acid positions randomized near the predicted stall site in the PTC, with only sequences containing stop codons shown. Log_2_-fold enrichment of sequences is plotted against total read count, for experiments carried out in duplicate. Blue, sequences enriched with an adjusted *P*-value < 0.01. **d**, Western blot of affinity-purified stalled CDH1-derived nascent chains. Reactions in the absence or presence of 50 µM PF846 are shown. A luciferase reporter with no PF846-dependent stalling sequence but with an N-terminal 3x-FLAG tag is shown as a control (lanes 1, 2). Lanes 3 and 4 are WT CDH1; lanes 5 and 6, HYHS-containing CDH1 sequence; lanes 7 and 8 are for CDH1-NPN*; lanes 9 and 10 are for CDH1-GCV*, * indicating the stop codon. Western blot of ribosomal protein RPS19 for each sample is also shown, as a loading control. **e**, Model of the mechanism of PF846-induced stalling. PF846 binding to the exit tunnel stalls the elongation of the nascent chains in the rotated state, with either A/A NC-tRNA or A/P NC-tRNA. Poor positioning of the 3’-CCA end prevents eEF2-GTP from catalyzing the translocation of hybrid A/P and P/E tRNAs into P/P tRNA and E/E tRNA sites. PF846 can also stabilize the resulting non-rotated ribosome with an empty A site that may signal quality control (QC) pathways.

Remarkably, we also identified many sequences with a stop codon at the final position enriched in a PF846-dependent manner in the PTC-proximal mRNA library (**Fig. 6c**), indicating that PF846 blocked proper translation termination of these NCs. These sequences, roughly 6-7% of the total termination sequences identified, have an entirely different amino acid consensus compared to those predicted to stall translation elongation (**Fig. 6a, 6c**, insets). Since PF846 and related compounds had not been observed previously to block translation termination, we tested whether PF846 affects termination at the identified sequences. Consistent with the enrichment observed from the PTC mRNA library, PF846 prevented NC release from ribosomes when sequences NPN, DPC, NVI and CVT–but not GCV–preceded the stop codon (**Fig. 6d**, **Extended Data 13d**). We did not observe an enrichment of eRF1 in the pelleted RNCs (**Extended Data 13d**), suggesting that aberrant termination induced by PF846 may prevent eRF1 binding.

### Model for PF846 induced translation stalling

The present structures of PF846-stalled CDH1-, PCSK9- and USO1-RNCs provide the first molecular detail for the selective stalling of nascent chains in human ribosomes by a small molecule. PF846 binds a newly identified small molecule-binding pocket in the exit tunnel of the human ribosome and traps select RNCs predominantly in the rotated state (**Fig. 6e**). The ability of PF846 to stall highly specific nascent chain sequences seems to require a distributed set of interactions between the nascent chain and the exit tunnel wall (**Fig. 4a**, **Extended Data Fig. 10**). The specific sequence determinants required for PF846-induced stalling of translation elongation extend roughly one-third of the way through the exit tunnel, beginning at the PTC and extending past the binding pocket for PF846 (**Extended Data Fig. 12**). Notably, sequences close to the PTC exert a more dominant effect on stalling potency, as revealed from the mRNA libraries (**Fig. 6**). However, sequences close to the PTC are not sufficient for stalling, as revealed in the Asn scans (**Extended Data Fig. 12**). In the case of PCSK9, the NC likely needs to span the length of the exit tunnel ^10^. The distributed nature of PF846-induced stalling is reminiscent of those observed in naturally-occurring stalling sequences in bacteria ^17,25^ and in eukaryotes ^26^. By contrast, the more general stalling induced by antibiotics bound to the bacterial ribosome exit tunnel may only require localized rearrangements between the drug binding site and the peptidyl transferase center ^27,28^.

Redirection of the nascent chain by PF846 seems to prevent stable base pairing of the NC-tRNA 3’-CCA end with the ribosomal P-loop, likely making these RNCs poor substrates for the GTPase eEF2 responsible for mRNA and tRNA translocation. Consistent with this model for stalling, mutations in the 3’-CCA end of A/P tRNA or P-loop of the bacterial ribosome that disrupt base pairing slow or prevent translocation ^29^. Furthermore, the SecM stalling peptide in bacteria is also capable of trapping the ribosome in the rotated state with hybrid A/P NC-tRNA, in which the 3’-CCA end of the tRNA is not properly base paired with the P-loop ^25^. Proper 3’-CCA end positioning has also been shown to be important for the peptidyl transferase reaction. Peptide bond formation is disrupted by certain amino acid sequences such as proline stretches that prevent pairing of the 3’-CCA end with the P-loop in ribosomal RNA. These sequences require dedicated translation elongation factors, EF-P in bacteria and eIF5A in eukaryotes, to rescue stalled ribosome by stabilizing the CCA end of the P-site tRNA in the conformation that favors peptide bond formation ^30^. Notably, although PF846 does not stall elongating ribosomes on NCs with proline sequences enriched near the PTC (**Fig. 6a**, inset), PF846 blocks termination when proline-rich sequences occur immediately adjacent to a stop codon positioned at the C-terminus of the CDH1 stalling sequence (**Fig. 6c**, inset). The broader sequence determinants for PF846-induced inhibition of termination remain to be elucidated, but may be distributed over a longer stretch of the NC, as seen for the hCMV stalling peptide, which ends in the sequence IPP but requires specific sequences in the NC that are span at least one-third the length of the exit tunnel ^18,31^. The mechanism of PF846-induced inhibition of translation termination differs from that of the hCMV peptide, which traps termination factor eRF1 on the ribosome in an inactive conformation ^18,31^. By contrast, the PF846-induced block of termination seems to have no observable effect on eRF1 binding (**Extended Data Fig. 13d**). This may be due to the PF846-stalled termination complex occupying the rotated state, thereby preventing eRF1 from accessing the tRNA A site on the small subunit, and/or preventing the 3’-CCA end of P-site tRNA from properly docking in the PTC ^32^.

Stalling of RNCs in the rotated state may serve as a critical determinant of PF846 selectivity by evading cellular quality control pathways that recognize aberrantly stalled ribosomes and initiate recycling ^33^. For example, Pelota (PELO, Dom34 in yeast) and its cofactors are involved in ribosome-associated quality control pathways that recognize stalled ribosome with an empty A site, to which Pelota binds and promotes ribosome subunit dissociation ^33^. Consistent with the hypothesis that quality control pathways resolve PF846-stalled RNCs, ribosome profiling experiments did not reveal a buildup of stalled RNCs in cells treated with PF846 ^10^. Furthermore, PELO and HBS1L are genetically linked to cellular toxicity of PF846-related compounds administered in high doses ^34^. The experimental methods presented here provide a means for probing the sequence specificity of PF846-like compounds, and should enable the discovery of other compounds that target the translating human ribosome for therapeutic effect.

## Supporting information

PF846 region coordinates and map from CDH1-RNC complex

## Acknowledgments

We thank Fuguo Jiang for his help in model building, Vignesh Kasinath, Junjie Liu, and Basil Greber for discussions about EM data processing and Kelly Nguyen for advice on signal subtraction methods, Meihua Tu and Nathanael Lintner for helpful suggestions. This work was funded by the Pfizer Emerging Science Fund, and by the NIH (R01-GM065050).

## Authors contributions

Conceptualization: KFM, SL, RD, and JHDC; Investigation and formal analysis: WL, FRW, KFM, STC, EM, and JHDC; Writing-original draft: WL and JHDC; Writing-review and editing: all authors.

## Author Information

RD is employed by Pfizer, Inc. KFM and SL were employed by Pfizer, Inc. The authors have filed patent applications related to this work.

## Methods

### DNA constructs and *in vitro* transcription

The DNA fragments needed to assemble the constructs encoding the affinity tagged nascent chains (EMCV IRES, CDH1 sequence encoding amino acids 586-750 and PCSK9 sequence encoding amino acids 1-35; **Fig. 1a**) were amplified from previous plasmids encoding full length PCSK9 and CDH1 ^10^. For the CDH1 nascent chain used for structural studies, we fused the EMCV IRES, affinity tags, CDH1 sequence encoding amino acids 586-750 and a NanoLuc luciferase reporter gene together by overlap PCR. For the PCSK9 nascent chain, CDH1 amino acids 716-750 were replaced with PCSK9 amino acids 1-35, leaving the CDH1-V domain intact. The USO1 construct was made by replacing the sequence encoding CDH1 amino acids 709-750 with the USO1 nascent chain sequence spanning that found previously to cause PF846-induced stalling (amino acids 258-299: RMKPWFEVGDENSGWSAQKVTNLHLMLQLVRVLVSPTNPPGA) ^10^. The NanoLuc control without stalling sequence was obtained by fusing the EMCV IRES and the NanoLuc open reading frame using overlap PCR. All PCR products were purified via spin columns (Qiagen) before their use in *in vitro* transcription reactions.

*In vitro* transcription reactions were performed using PCR products generated with primers encoding a flanking T7 RNA polymerase promoter and a poly-A tail. Reactions were set up with 20 mM Tris-HCl pH 7.5, 35 mM MgCl_2_, 2 mM spermidine, 10 mM DTT, 1 U/mL pyrophosphatase (Sigma), 7.5 mM each NTP, 0.2 U/L SUPERaseIn RNase Inhibitor (ThermoFisher), 0.1 mg/mL T7 RNA polymerase and 40 ng/µL DNA. After 3 hours incubation at 37 °C, 0.1 U/µL DNase I (Promega) was added to the reactions, and incubated at 37 °C for 30 min to remove the template DNA. RNA was precipitated for 2-3 hours at –20 °C after adding 1/2 volume of 7.5 M LiCl/50 mM EDTA, and the resulting pellet was washed with cold 70% ethanol and dissolved with RNase free water. The mRNA was further purified by Zymo RNA Clean and Concentrator (Zymo research) before use in *in vitro* translation reactions.

### *In vitro* translation reactions

The HeLa cell extract was made as described previously ^10^. Briefly, a frozen HeLa cell pellet was thawed and suspended with an equal volume of lysis buffer (20 mM HEPES pH 7.5, 10 mM potassium acetate, 1.8 mM magnesium acetate, and 1 mM DTT). After incubation on ice for 20 min, the cells were lysed with a Dounce homogenizer 150 times, followed by centrifugation twice at 1,200 x *g* for 5 min. The supernatant after centrifugation was aliquoted to avoid freeze-thaw cycles, and flash-frozen in liquid nitrogen. For a 10 µL reaction, 5 µL HeLa cell extract was used in a buffer containing final concentrations of 20 mM Hepes pH 7.4, 120 mM KOAc, 2.5 mM Mg(OAc)_2_, 1 mM ATP/GTP, 2 mM creatine phosphate (Roche), 10 ng/µL creatine kinase (Roche), 0.21 mM spermidine, 0.6 mM putrescine, 2 mM TCEP, 10 µM amino acids mixture (Promega), 1 U/µL murine RNase inhibitor (NEB), 200 ng mRNA, and 0.1 mM PF846 in 1% DMSO, or 1% DMSO as control. Translation reactions were incubated for 23 min at 30 °C, after which luciferase activity was monitored with a Microplate Luminometer (Veritas). Before preparation of PF846-dependent stalled RNCs, all chimeric mRNAs were tested in in vitro translation reactions for PF846-induced stalling, using reactions plus or minus 50 µM PF846.

### Purification of stalled RNCs

1.5 mL *in vitro* translation reactions with 0.2 mM PF846 were incubated at 30 °C for 23 min and then centrifuged at 11,400 r.p.m. for 5 min. The supernatant was incubated with 50 µL (packed volume) of anti-FLAG M2 agarose beads (Sigma) for 30 min at room temperature with gentle mixing. To avoid the binding of nonspecific ribosomes and other proteins, the beads were washed at room temperature three times with 500 µL RNC buffer (20 mM Hepes pH 7.4, 300 mM potassium acetate, 5 mM magnesium acetate, 1 mM DTT, 0.2 mM PF846), then three times with 500 µL RNC buffer plus 0.1% TritonX-100, followed by three times with 300 µL RNC buffer plus 0.5% TritonX-100, and finally washed twice with 300 µL RNC buffer. The PF846 stalled RNCs, bound to the FLAG beads by the N-terminal 3xFLAG tag, were eluted twice at room temperature for 20 min each time, with 30 µL 0.2 mg/mL 3xFLAG peptide (Sigma) in RNC buffer. The eluted fractions were combined and loaded onto a 50% sucrose cushion prepared with cushion buffer (25 mM HEPES-KOH pH 7.5, 120 mM KOAc and 2.5 mM Mg(OAc)_2_, 1 M sucrose, 1 mM DTT, 0.2 mM PF846), and centrifuged for 1 hour in an MLA rotor (Beckman Coulter) with 100,000 r.p.m. (∼603,000 *g*) at 4 °C. The pellet was suspended in ice-cold RNC buffer and was immediately used for cryo-EM grid preparation. The concentration of purified RNC was determined using a NanoDrop Microvolume Spectrophotometer and calculated using 1 A_260_ unit corresponding to 20 pmol of ribosome.

### RNaseA digestion and Western blot

RNCs released from the anti-FLAG beads and pelleted through the sucrose cushion were resuspended in 20 µL to an A_260_ O.D. of around 1 and then incubated with or without 100 µg/mL DNase free Rnase A (ThermoFisher) and 50 mM EDTA at 37 °C for 30 min followed by Western blot of the FLAG-tagged peptides. For the Western blots, Monoclonal ANTI-FLAG^®^ M2-Peroxidase (HRP) antibody (SIGMA) at 1:10,000 dilution was used.

### Cryo-sample preparation and Data collection

Approximately 3 µL of freshly made RNC at a concentration of 40 to 60 nM were incubated for 1 min on plasma-cleaned 300-mesh holey carbon grids (C-flat R2/2), on which a home-made continuous carbon film was pre-deposited. Grids were blotted for 4 seconds with 100% humidity at 4 °C and plunge-frozen in liquid ethane using an FEI Vitrobot. Automated data collection was performed on a Titan Krios electron microscope (FEI) equipped with a K2 Summit direct detector and GIF Quantum filter (Gatan) at 300 kV (**Extended Data Tables 1-4**). The total exposure time was 9 seconds, with a total dose of 50 electrons per Å^2^ (frame dose 1.3 electrons per Å^2^).

### Data Processing

Frames were aligned using MotionCor2 with FtBin 2 ^36^. We used Gautomatch (http://www.mrc-lmb.cam.ac.uk/kzhang/) for automatic particle picking. Power spectra, defocus values and astigmatism were determined with Gctf software with per-particle CTF estimation ^37^. All the subsequent data processing was performed in RELION 2.0 ^38^ and RELION 3.0 ^39^ unless specifically noted. To speed up computing, we binned the particles by 8 times during particle extraction. Reference free 2D classification was performed to remove particles with ice or other contaminants, followed by 3D classification to select RNC particles. The initial 3D classification roughly separated the ribosome particles from the non-ribosome particles without yielding structurally distinct classes.

For the CDH1-RNC data set, we divided the whole data set into three subsets and carried out particle sorting as shown in **Extended Data Fig. 2a**. After initial 3D classification and refinement, an additional round of 3D classification was performed with a local angular search that separated CDH1-RNCs in rotated states from a class with apo-80S ribosomes or ribosomes with weak density for tRNA (**Extended Data Fig. 2a**). A total number of 84,437 particles representing the CDH1-RNC in the rotated state were combined and subjected to 3D refinement, which generate a map of 4.0 Å without the use of a mask. Signal subtraction was conducted to improve the density of the nascent peptide chain ^40^. Starting with the aligned 84,437 particles, a region with CDH1 nascent chain, PF846 and A/P NC-tRNA was extracted and classified by applying a soft mask generated in RELION. Focused 3D classification separated the rotated state into two major classes, one with A/P NC-tRNA and P/E tRNA (84% of particles), and the other with A/A NC-tRNA and P/E tRNA (15% of the particles). The more-abundant class with A/P NC-tRNA had good electron density for the CDH1 nascent chain, which was used for model building (**Extended Data Fig. 2a**). The overall resolutions of the CDH1-RNC’s with A/P NC-tRNA and A/A NC-tRNA were 3.0 Å and 3.9 Å, respectively, using the gold-standard Fourier shell correlation (FSC=0.143) criterion (**Extended Data Fig. 2b**, **Extended Data Table 1**).

To identify other CDH1-RNC classes, we combined the remaining classes after sorting the RNC with hybrid tRNAs (**Extended Data Fig. 2a**) for an additional 3D refinement and classification with a local angular search, from which three major classes were observed: 80S ribosome with GTPase bound in the A site (24%), CDH1-RNC with P/P tRNA (26%), and the large (60S) subunit (15%). Another round of signal subtraction for the CDH1-RNC with P/P tRNA was conducted based on the P/P tRNA density, resulting in ∼61% particles which were chosen for the final reconstruction of the non-rotated state RNC.

Two data sets for the PCSK9-RNC were collected on the same Titan Krios microscope within a week of each other. After overall refinement, an additional round of 3D classification was performed with a local angular research, which separated RNCs in the rotated state (**Extended Data Fig. 3a**). We combined a total of 53,247 particles with hybrid state tRNAs from both data sets for another round of 3D auto refinement. The post-processing step implemented in RELION 3.0 was used to generate the final maps using the FSC=0.143 criterion indicating an average resolution of 3.0 Å (**Extended Data Fig. 3b, Extended Data Table 2**) ^41,42^. Signal subtraction for PCSK9-RNC with either hybrid tRNAs or P/P tRNA was conducted in the same way as described for the CDH1-RNC complexes ^40^ (**Extended Data Fig. 3**). Data for the USO1-RNC were collected in a single session and processed similarly to the PCSK-RNC sample, except signal subtraction yielded only a single tRNA population, with A/P NC-tRNA (**Extended Data Fig. 4**, **Extended Data Table 4**).

For the data set of stalled PCSK9-RNC prepared using a short time interval, the selected particles after RELION 2D classification were processed using CryoSPARC ^43^ for further 3D classification and homogeneous refinement, which generated two maps representing different states of ribosome at an average resolution of 4.6 Å and 4.8 Å respectively according to the “gold standard” FSC=0.143 (**Extended Data Fig. 3d, e, Extended Data Table 3**) ^41,44^.

### Model Building and Refinement

Initial rigid body fitting of the human 40S ribosomal subunit structure (PDB: 6ek0) 45 and 60S subunit structure (PDB: 5aj0) ^46^ was done for all the rotated RNCs (CHD1-RNCs and PCSK9-RNCs with either A/A NC-tRNA and A/P NC-tRNA, and USO1-RNCs with A/P NC-tRNA), using UCSF Chimera ^47^. We then docked the ribosomal proteins and RNAs of the 40S and 60S subunits separately into the EM map, with parts of the model manually adjusted to the EM map using COOT ^48^. For rotated-state RNCs, helix H69 in 28S rRNA was modeled based on the rotated state structure in PDB entry 4u4r ^7^, followed by manual rebuilding in COOT. All the tRNAs (A/A, A/P, P/E tRNA) for the RNCs in the rotated state were docked into the EM maps using reported tRNA structures (PDB: 3j7r) ^49^, followed by manual rebuilding in COOT ^48^. The anticodon nucleotides from 34 to 36 in the tRNA were modeled as one sequence (CCG). RNCs with P/P site tRNA were initially modeled using rigid-body docking of the human ribosome structure (PDB: 5aj0) ^46^ and then corrected based on the EM density in COOT. The nascent chains for the CDH1-RNC and PCSK9-RNC were manually built into the density in one sequence register (CDH1_NC sequence: CRKAQPVEAGLQIPAILGILGGILALLILILLLLLF and PCSK9_NC sequence: WWPLPLLLLLLLLLGPAGARAQED) in COOT. The USO1-RNC NC was modeled as poly-alanine, due to the lack of recognizable density features for side chains. All RNC structures were refined in PHENIX using phenix.real_space_refine with secondary structure restraints imposed ^50^. Model refinement and validation statistics are provided in **Extended Data Tables 1–4**.

To model the interaction of *E. coli* 23S rRNA and PF846, the high-resolution model (PDB: 4ybb) ^51^ was docked into the map by rigid-body docking. For the analysis of the pairing between the A/P-tRNA 3’-CCA end and 28S rRNA, we docked the high resolution tRNA (PDB: 1vy4) ^35^ as a rigid body, using nucleotides in the ribosomal P-loop as reference. Subsequent modeling of C75 used manual rebuilding in Coot followed by real-space refinement, enforcing minimal changes to the positions of C74 and A76.

### Modeling of codon-anticodon base pairs

Correlations between modeled nucleotides and observed cryo-EM density at 3.0 Å resolution are dominated by the phosphate-ribose backbone and the common position of the nucleobase in pyrimidines and five-membered ring in purines. Thus, it is difficult to distinguish purines from pyrimidines at this resolution using only real-space correlations. We therefore used helix h24 in human 18S rRNA (nts 1037-1078) to assess the ability of real-space difference density maps to distinguish purines and pyrimidines in base pairs, and guanosines from adenosines in base pairs, in ∼3.0 Å resolution cryo-EM maps. We generated map fragments from the CDH1-RNC cryo-EM map with A/P NC-tRNA and made mutations in nucleotides using COOT ^48^, followed by real-space refinement of models in Phenix (phenix.real_space_refine) ^50^, with secondary structure restraints generated by phenix.secondary_structure_restraints and verified manually. We then generated real-space difference maps in Chimera ^47^, as follows. We set the minimum threshold of the observed cryo-EM map to zero (vop threshold minimum 0.), and used the model coordinates to generate a calculated map at 3.0 Å resolution (molmap). Then, we generated (observed – calculated) difference maps by minimizing the root-mean-square of the differences (vop subtract minRMS true). From these maps, we found that negative difference density was diagnostic for positions that are incorrectly modeled as purine, due to negative density on the N2, C2, N1, C6, and O6/N6 positions of the modeled purine ring. Notably, we also observed positive difference density in these maps consistent with the position of a purine incorrectly modeled as a pyrimidine, but not reproducibly. This is possibly due to the distribution in density in the map calculated from model coordinates with ADP values grouped at the nucleotide level, currently the only setting available in phenix.real_space_refine ^50^.

We observed that, in the CDH1-RNC and PCSK9-RNC reconstructions at ∼3.0 Å resolution, the cryo-EM density between the nucleobases for base pairs in 18S rRNA was clearly separated, and could be easily assigned to purines versus pyrimidines visually. However, the density between the nucleobases in the tRNA anticodon-mRNA codon base pairs was fused. Furthermore, the density shapes did not clearly align with either a pyrimidine or purine base, as shown in **Fig. 2** and **Extended Data Fig. 6**.

### Quantum mechanical calculations of PF846 conformations

All torsional assessments and geometry optimizations of PF846 were performed by quantum mechanical calculations using TeraChem ^52,53^ using a DFT-(b3lyp) level of theory and a 6-31gs basis set. Dihedral angle energy profiles from model systems derived from truncation of the small molecule X-ray structure of PF846 were used to create low energy poses for fitting to the density. All poses were geometry optimized before fitting. **Extended Data Table 5** shows the calculated energies for the poses in **Extended Data Fig. 6**. After rigid-body docking of PF846 pose 4, the torsion angles between the chloropyridine and amide linkage, and between the aryl ring and amide linkage were manually adjusted by ∼15° each to better fit the map density.

### DNA libraries with randomized codons in the CDH1 stalling sequence

DNA libraries encoding the open reading frame containing the CDH1 stalling sequence in **Fig. 1a**, with stretches of 4 random codons encoded by NNK sequences (N=any nucleotide, K=G or T) to introduce random mutations in different sites of the CDH1 nascent chain within the ribosome exit tunnel, were generated by PCR amplification. PCR products were purified using spin columns (Qiagen, catalog number: 28506), then treated with Dpn1 nuclease at 37 °C for 1 hour to remove template DNA. A double-stranded DNA fragment encoding a T7 RNA polymerase promoter was then ligated to the 5’ end of the treated DNA, along with a double-stranded DNA encoding a poly-A tail to the 3’ end, using T4 DNA ligase and T4 Polynucleotide Kinase at 16 °C overnight. The ligation reaction was followed by PCR amplification using primers flanking the T7 promoter and poly-A tail to generate the final DNA library.

The following primers were used to make the NNK DNA libraries.

Lib_PTC (nucleotides corresponding to the amino acids near the peptidyl transferase center): 5’-NNKNNKNNKtttcttcggaggagagcggtgg-3’ and 5’-MNNcagaatcagaattagcaaagcaagaattcctcc-3’; Lib_compound (nucleotides corresponding to the amino acids predicted to interact with the compound in the stalled complex): 5’-NNKNNKNNKattctgctgctcttgctgtttcttcg-3’ and 5’-MNNagcaagaattcctccaagaatcccc-3’; In these libraries, M stands for A or C nucleotides.

### Selection of ribosome nascent chain complexes stalled by PF846 using mRNAs encoding random libraries

The mRNAs encoding the NNK libraries were prepared by T7 RNA polymerase transcription of the above NNK DNA libraries, followed by mRNA purification as described above. The mRNA libraries were incubated in a 1.0 mL *in vitro* translation reaction with 50 µM PF-06446846 at 30 °C for 28 min. Stalled RNCs were then purified using anti-FLAG peptide beads as described above, with the following modifications. Stalled ribosome nascent-chain complexes (RNCs) were bound to the beads in the presence of 1 µM DNA oligonucleotide 5’-TCTCCTCCGAAGAAA-3’ (targeting DNA) and 2 µL RNase H (5 units/µL, NEB) were added to the reaction. The targeting DNA was designed to base pair with the CDH1 mRNA codons protected in the mRNA binding channel of stalled RNCs, but that would be exposed in free mRNAs, or in mRNAs bound to elongating ribosome complexes. RNCs were then eluted from the anti-FLAG peptide beads using the competitor FLAG peptide, and pelleted through a sucrose cushion as described above. After the sucrose cushion pelleting step, the pellet was resuspended in 40 µL RNase H buffer (20 mM Tris 7.5, 150 mM KCl, 2.5 mM MgCl_2_, 2mM DTT) and treated with 1 µL (5 units) RNase H and 1 µM of the targeting DNA oligonucleotide at 37 °C for 40 min.

Total RNA was then extracted from the purified and RNase H treated RNCs using TRIzol LS (Thermo Fisher Scientific, cat. number: 10296010). First strand cDNAs were synthesized using a primer specific to a nucleotide sequence in the open reading frame of **Fig. 1a** located 3’ of the stalling sequence, using Superscript II (Invitrogen, cat. number: 18064014), according to the manufacturer’s protocol. The cDNA was then used in a 70 µL PCR reaction with Q5 DNA polymerase with optimized cycles of 10 s at 98 °C, 30 s at 66 °C and 15 s at 72 °C, followed by a two step purification with SPRIselect beads (Beckman Coulter, cat. number: B23317), first by adding 42 µL beads (0.6X amount of the PCR reaction) and then applying the flowthrough to another 56 µL beads (0.8X amount of the PCR reaction). The resulting bound DNA was eluted in 15 µL molecular biology grade water. The DNA libraries were sequenced using an Illumina HiSeq 4000. The following oligonucleotides were used to prepare the sequencing libraries.

RT primer used for synthesizing the first strand cDNA, and containing a 10-nt unique molecular identifier (UMI) barcode: 5’-AGACGTGTGCTCTTCCGATCT(N)_10_CTCTTTGACCACCGCTCTCC-3’ Universal P5 adaptor (Forward primer for PCR): 5′-AATGATACGGCGACCACCGAGATCTACACACACTCTTTCCCTACACGACGCTCTTC CGATCTNNNNNCGAAGCAGGATTGCAAATTCCT-3’ Indexed P7 adapter (Reverse primer for PCR): 5’-CAAGCAGAAGACGGCATACGAGATCCTTGGAAGTGACTGGAGTTCAGACGTGTGC TCTTCCGATCT-3’ (the underlined nucleotides indicate the index sequence for one sample. Different index sequences were used for every sample).

### Analysis of the Illumina sequencing libraries

UMIs were extracted from raw reads using umi_tools ^54^ and sequences were aligned to an index of all 12-mer DNA sequences using seqkit ^55^ and bowtie2 ^56^ (parameters --N 0, --no-1mm-upfront, -norc --L 24). No mismatches were allowed in the 6 bp sequences flanking the library bases to ensure each read was a correctly generated library member. Alignments were sorted and indexed with samtools ^57^ and deduplicated with umi_tools using the ‘unique’ method. Deduplicated reads were tabulated by translated library sequence, replicate, and condition (PF846 treated or untreated) in python, and PF846 sensitivity was analyzed using DESeq2 ^58^. PF846-sensitive or -resistant library members were called with a false discovery rate cutoff of 1%. Sequencing data after analysis is included in **Extended Data Table 6**.

### Validation of PF846-dependent stalling sequences and inhibition of termination

To validate the sequences identified in the mRNA selections are bona-fide targets of PF846-dependent stalling, we cloned reporter mRNAs with representative examples into the CDH1-chimeric mRNA construct described above and shown in **Fig. 1a**. We replaced codons for amino acids L726-L729 in the CDH1 sequence with codons for the following sequences: HYHS and RSCK for sequences stalled by PF846 during elongation, and NPN*, CVT*, DPC*, and NVI* for sequences that lead to PF846-dependent inhibition of termination (*, stop codon). We also used GCV*, which was not enriched in the experiment, as a negative control for PF846-dependent inhibition of translation termination. Codons used for each sequence are the following:

HYHS–CACTACCACTCT

RCSK–CGATCTTGTAAG

NPN*–AATCCAAACTAA

CVT*–TGTGTCACATAA

DPC*–GATCCTTGTTAG

NVI*–AATGTTATTTAG

As an additional negative control, we cloned nanoluciferase with an N-terminal 3X-FLAG tag.

To test whether the sequences HYHS and RSCK, and the ones with stop codons, lead to PF846-dependent stalling in the context of the CDH1 sequence, we synthesized mRNAs as described above and subjected them to *in vitro* translation reactions, also as described above, with the following modification. We carried out the *in vitro* translation reactions for nascent chains with HYHS and RSCK sequences using different concentration of PF846, in order to determine the IC50 values for PF846 for each sequence, in comparison to that for the WT CDH1 sequence. To assess the amount of NCs that remain associated with ribosomes after the reactions, we also carried out *in vitro* translation reactions with or without 50 µM PF846, followed by pelleting ribosomes through a sucrose cushion and Western blotting with an anti-FLAG antibody (Monoclonal ANTI-FLAG^®^ M2-Peroxidase (HRP) antibody (SIGMA)) and RPS19 antibody (Bethyl Laboratories) as a loading control. We also probed for the presence of eRF1 using Western blotting after the ribosome pelleting step (anti-eRF1 antibody from Cell Signaling Technology).

### Figure preparation

Figures were generated with UCSF Chimera ^47^, PyMOL (Schrödinger) and ChemDraw (PerkinElmer).

### Data availability

The maps have been deposited with the Electron Microscopy Data Bank under the accession codes EMD-— Atomic Coordinates and structure factors have been deposited in the Protein Data Bank with accession codes —.

