## Supplementary Figures and Tables for "Structural basis for selective stalling of human ribosome nascent chain complexes by a drug-like molecule"

### Extended Data Figures and Tables

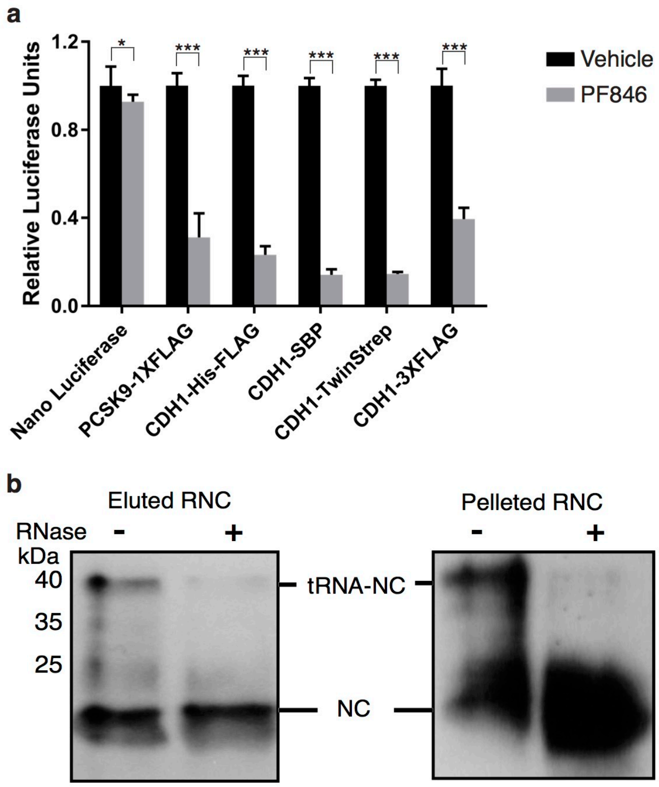

**Extended Data Fig. 1 | PF846-mediated stalling of RNCs. a**, *In vitro* translation assays of PCSK9 and CDH1 nascent chains with different affinity tags in the absence (black bar) or presence (grey bar) of 100 μM PF846. Significance value: * p<0.1, **p<0.001. **b**, Western blot with anti-FLAG antibody, plus or minus RNase A treatment of affinity purified stalled PCSK9-RNCs after elution and pelleting through a sucrose cushion.

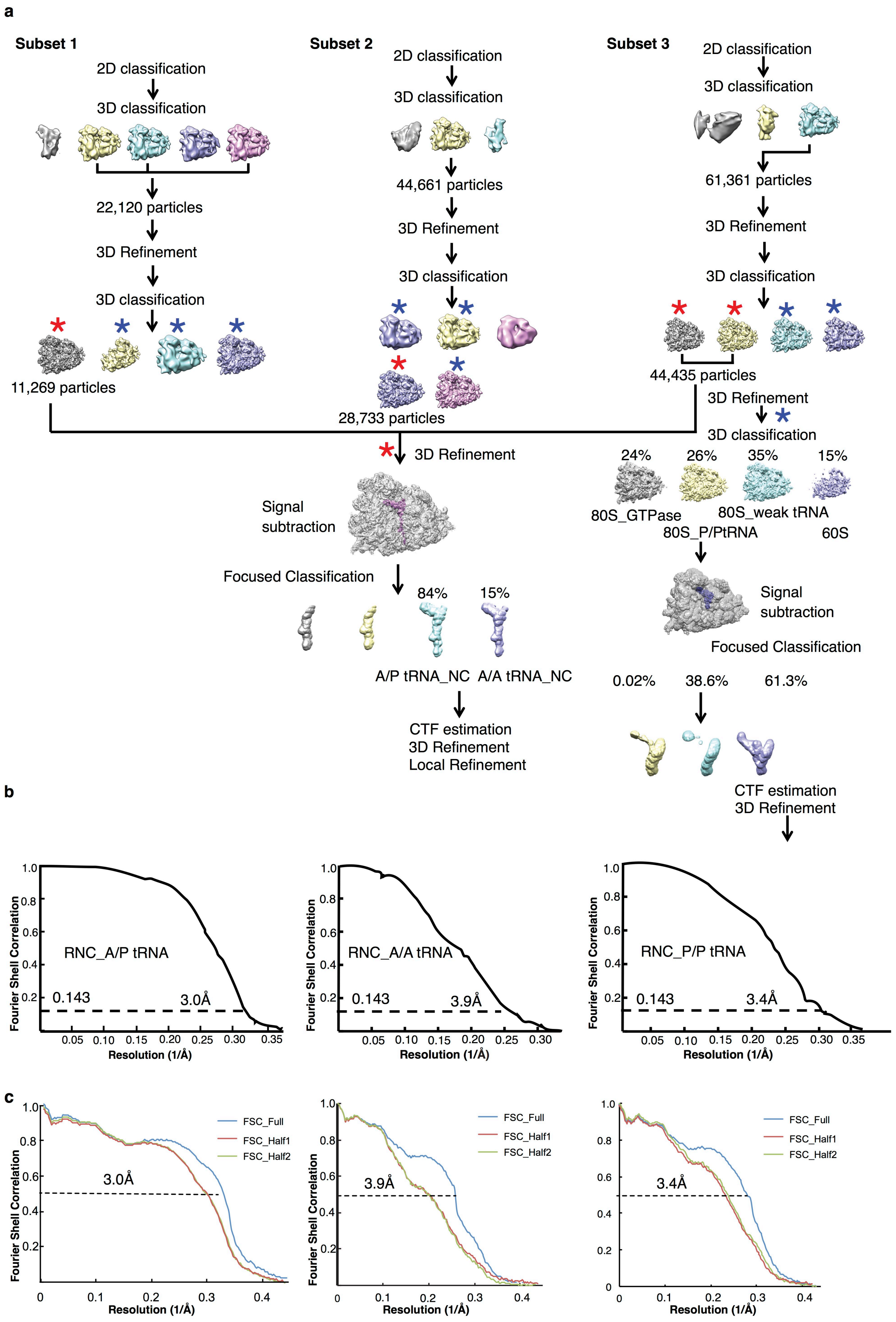
a
Subset 1
2D classification
3D classification
22,120 particles
3D Refinement
3D classification
11,269 particles
Subset 2
2D classification
3D classification
44,661 particles
3D Refinement
3D classification
28,733 particles
Subset 3
2D classification
3D classification
61,361 particles
3D Refinement
3D classification
44,435 particles
3D Refinement
3D classification
24%
26%
35%
15%
80S_GTPase
80S_weak tRNA
80S_P/PtRNA
60S
3D Refinement
Signal subtraction
Focused Classification
84%
15%
A/P tRNA_NC
A/A tRNA_NC
CTF estimation
3D Refinement
Local Refinement
Signal subtraction
Focused Classification
0.02%
38.6%
61.3%
CTF estimation
3D Refinement
b
RNC_A/P tRNA
0.143
3.0Å
RNC_A/A tRNA
0.143
3.9Å
RNC_P/P tRNA
0.143
3.4Å
Fourier Shell Correlation
Resolution (1/Å)
c
FSC_Full
FSC_Half1
FSC_Half2
3.0Å
3.9Å
3.4Å
Fourier Shell Correlation
Resolution (1/Å)

**Extended Data Fig. 2 | Cryo-EM data processing and model validation of PF846 stalled CDH1-RNCs. a**, The EM micrographs were first divided into 3 subsets for classification and refinement. The selected classes after refinement (labeled with red asterisks) were combined for an overall refinement. Signal subtraction using the A/P-tRNA, nascent chain and PF846 allowed classification and refinement of separate classes with A/A NC-tRNA or A/P NC-tRNA. The rest of the particles remaining after selecting the rotated state RNCs (labeled with blue asterisks) were merged and used for another refinement and 3D classification, which generated 4 different classes, including a non-rotated state with P/P NC-tRNA. **b**, Final FSC curves of CDH1_RNCs, with the “gold standard” value of 0.143 used to define the resolution indicated. **c**, Model to map correlation with resolution at the FSC value of 0.5 shown.

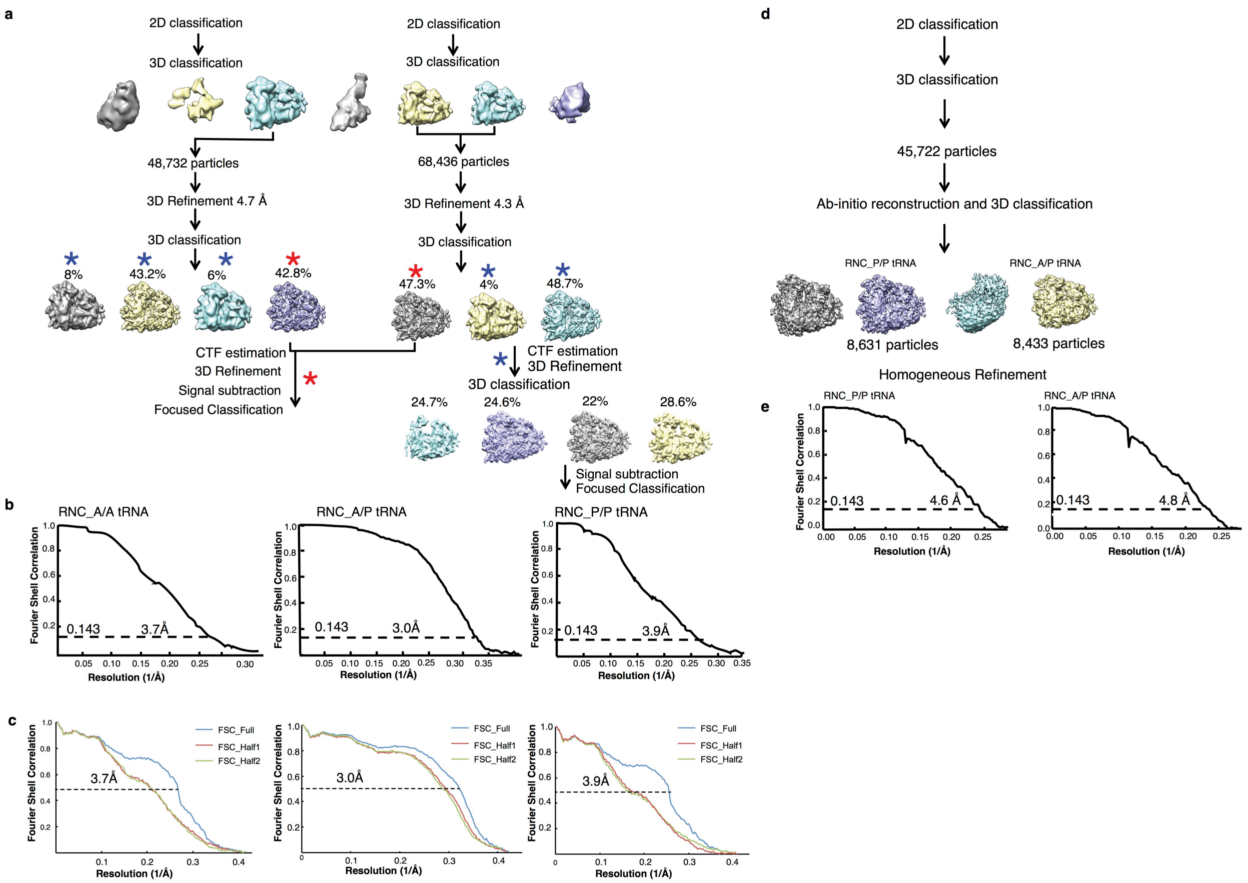

**Extended Data Fig. 3 | Cryo-EM data processing of PF846 stalled PCSK9-RNCs. a,** Data processing steps for stalled PCSK9-RNC, as described in **Extended Data Fig. 2**. **b,** Final FSC curves of PCSK9_RNCs, with the "gold standard" value of 0.143 used to define the resolution indicated. **c,** Model to map correlations with resolution at FSC value of 0.5. **d,** Data processing steps for sample prepared with a short incubation time, which generated a higher ratio of non-rotated to rotated states of the ribosome. **e,** Final FSC curves of RNCs, with the "gold standard" value of 0.143 used to define the resolution as indicated.

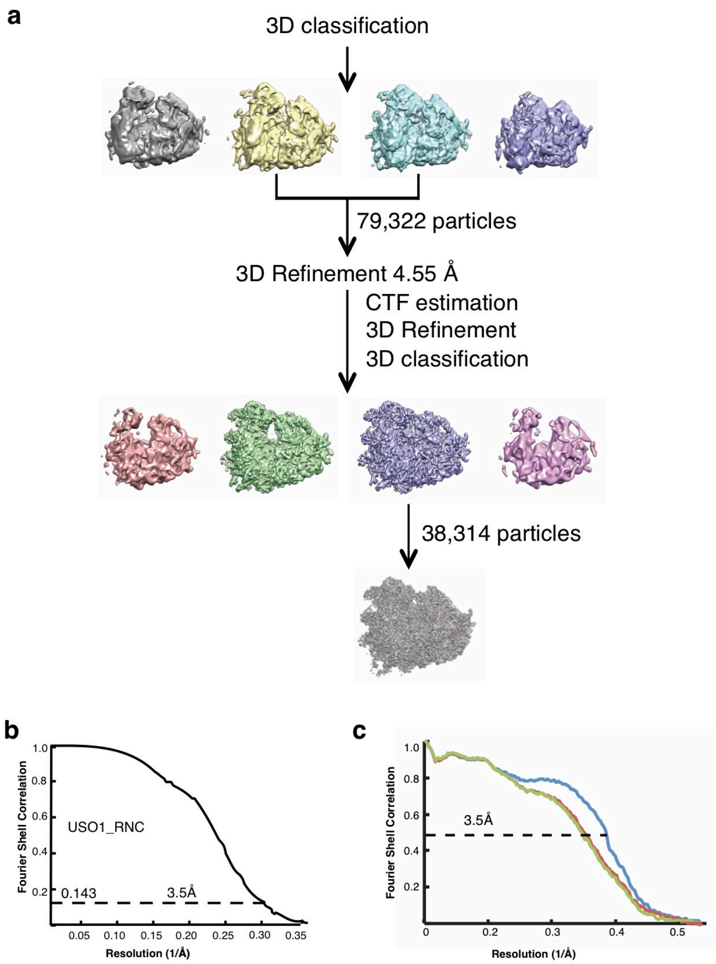

**Extended Data Fig. 4 | Cryo-EM data processing and model validation of PF846 stalled USO1-RNCs. a**, Data processing steps for USO1-RNC complexes. **b**, Final FSC curve of USO1-RNC, with the “gold standard” value of 0.143 used to define the resolution indicated. **c**, Model to map correlation with the resolution at an FSC value of 0.5.

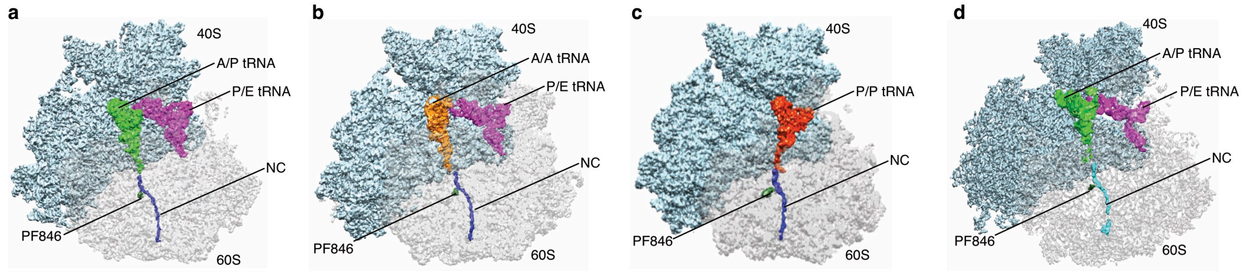

**Extended Data Fig. 5 | Cryo-EM reconstructions of PF846-stalled PCSK9 and USO1 RNCs. a,** PCSK9-RNC in the rotated state with A/P and P/E tRNA, with 40S subunit (light blue), 60S subunit (grey), A/P site tRNA (green), P/E site tRNA (magenta) stalled PCSK9 nascent chain (blue) and PF846 (dark green) shown. **b,** PCSK9-RNC in the rotated state with A/A tRNA (orange) and P/E tRNA (magenta). **c,** Non-rotated state of the PCSK9-RNC stalled by PF846, with P/P tRNA (dark orange). **d,** USO1-RNC in the rotated state with A/P and P/E tRNA. Color coding as in panel (**a**), but with the NC in cyan.

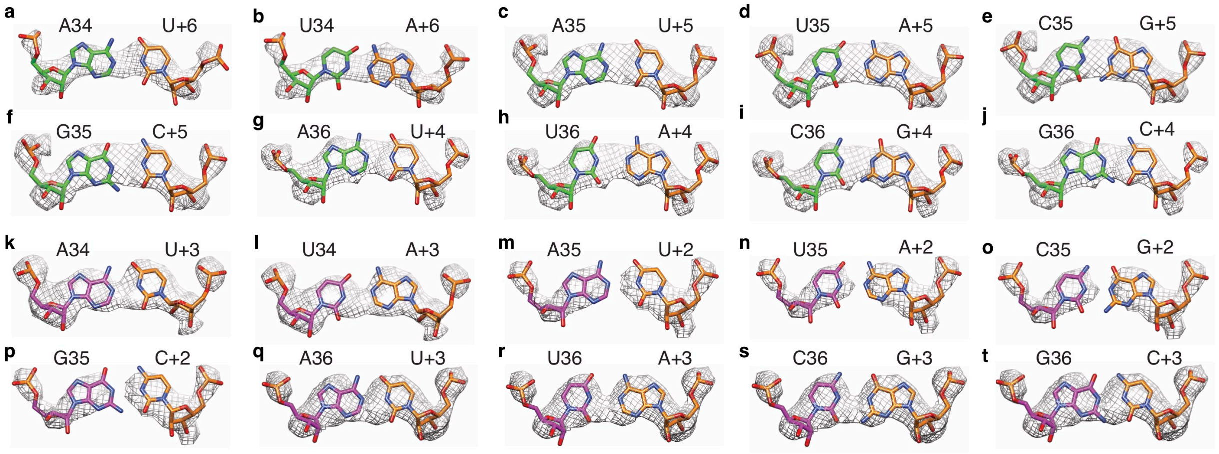

**Extended Data Fig. 6 | Cryo-EM density maps for tRNA-mRNA base pairs in the CDH1-RNC complex in the rotated state with A/P and P/E tRNAs. a-b**, Models for an A-U or U-A base pair between A/P tRNA anticodon nucleotide 34 (green) and mRNA codon nucleotide +6 (3rd nucleotide of codon in the A site)(orange). **c-f**, Models for A-U, U-A, G-C or C-G base pairs between A/P tRNA anticodon nucleotide 35 and mRNA codon nucleotide +5 (2nd nucleotide of codon in the A site). **g-j**, Models for A-U, U-A, G-C or C-G base pairs between A/P tRNA anticodon nucleotide 36 and mRNA codon nucleotide +4 (1st nucleotide of codon in the A site). **k-l**, Models for an A-U or U-A base pair between P/E tRNA anticodon nucleotide 34 (magenta) and mRNA codon nucleotide +3 (3rd nucleotide of codon in the P site)(orange). **m-p**, Models for A-U, U-A, G-C or C-G base pairs between P/E tRNA anticodon nucleotide 35 and mRNA codon nucleotide +2 (2nd nucleotide of codon in the P site). **q-t**, Models for A-U, U-A, G-C or C-G base pairs between P/E tRNA anticodon nucleotide 36 and mRNA codon nucleotide +1 (1st nucleotide of codon in the P site). All models are shown refined into the observed CDH1-RNC density.

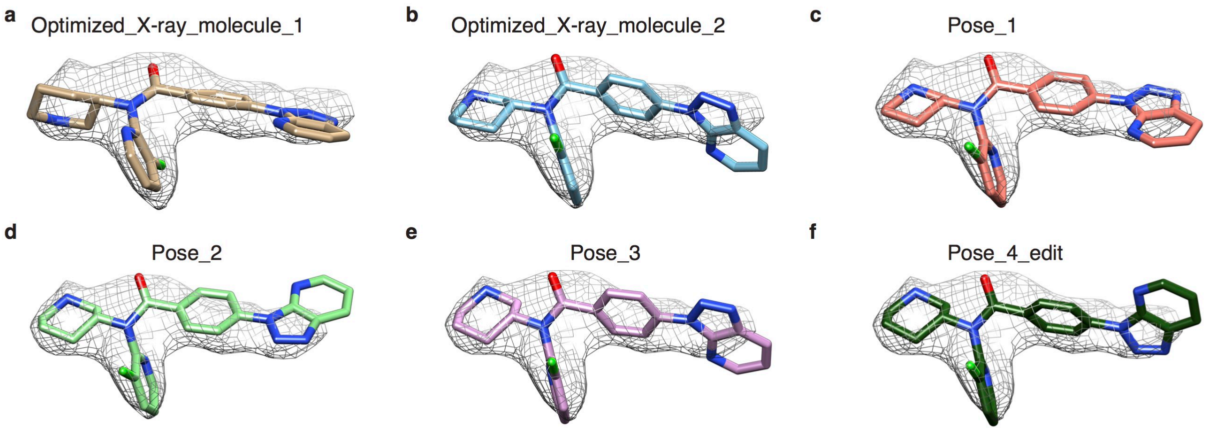

**Extended Data Fig. 7 | Docking of different conformations of PF846 into density extracted from the CDH1-RNC cryo-EM map. a-b,** X-ray structures of PF846, with (**a**) as molecule 1 and (**b**) as molecule 2 [10]. **c-f,** Low energy conformations of PF846 based on quantum mechanical calculations starting from the X-ray structures of molecule 2. From **c** to **f** are poses 1 to 4 in **Extended Data Table 5**. Note that pose 4 was manually adjusted to fit the density by ~15° dihedral rotations of the chloropyridine and benzyl rings.

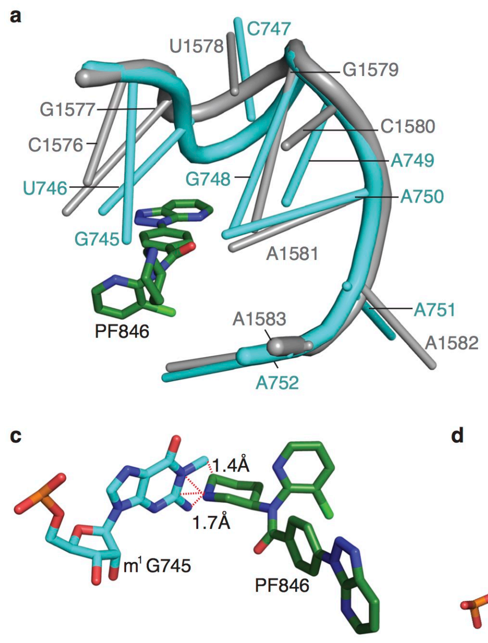

**Extended Data Fig. 8 | Modeling of PF846 in the predicted binding pocket of *E. coli* ribosome rRNA. a,** Cartoon representation of the conserved small molecule binding loop from *E. coli* and human ribosomes, colored in dark cyan and gray, respectively. **b,** Residues that contribute to the binding of PF846 are shown in stick format (PDB: 4ybb). **c-e,** 16S rRNA residues from *E. coli*, which would have direct interactions with PF846 based on the corresponding positions in the human ribosome, and would lead to steric clashes. Dashed lines in **c-e** indicate the closest distances between the nucleotides and PF846. In *E. coli*, G745 is modified to *N1*-methyl-G, and U746 to ψ746, although the steric clashes would occur even with unmodified nucleotides.

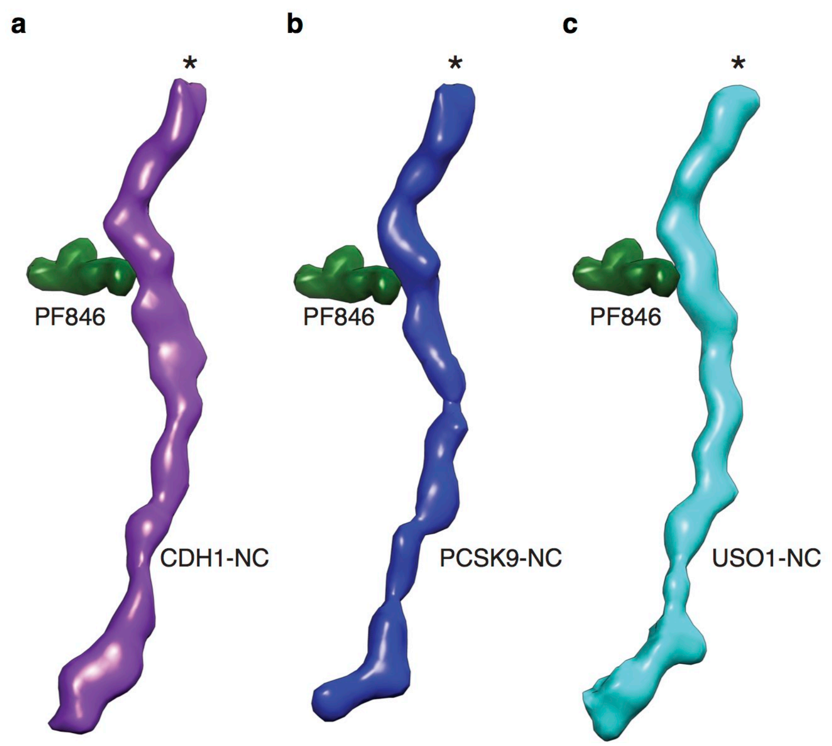

**Extended Data Fig. 9 | Cryo-EM densities of PF846 stalled nascent chains.** Density extracted from maps of the **a**, CDH1-RNC, **b**, PCSK9-RNC and **c**, USO1-RNC in the rotated state with A/P NC-tRNA, shown in the same orientation. The maps were low-pass filtered to 4.5 Å, with the asterisks indicating the location of the peptidyl transferase center. The orientation is similar to that in **Fig. 1c**.

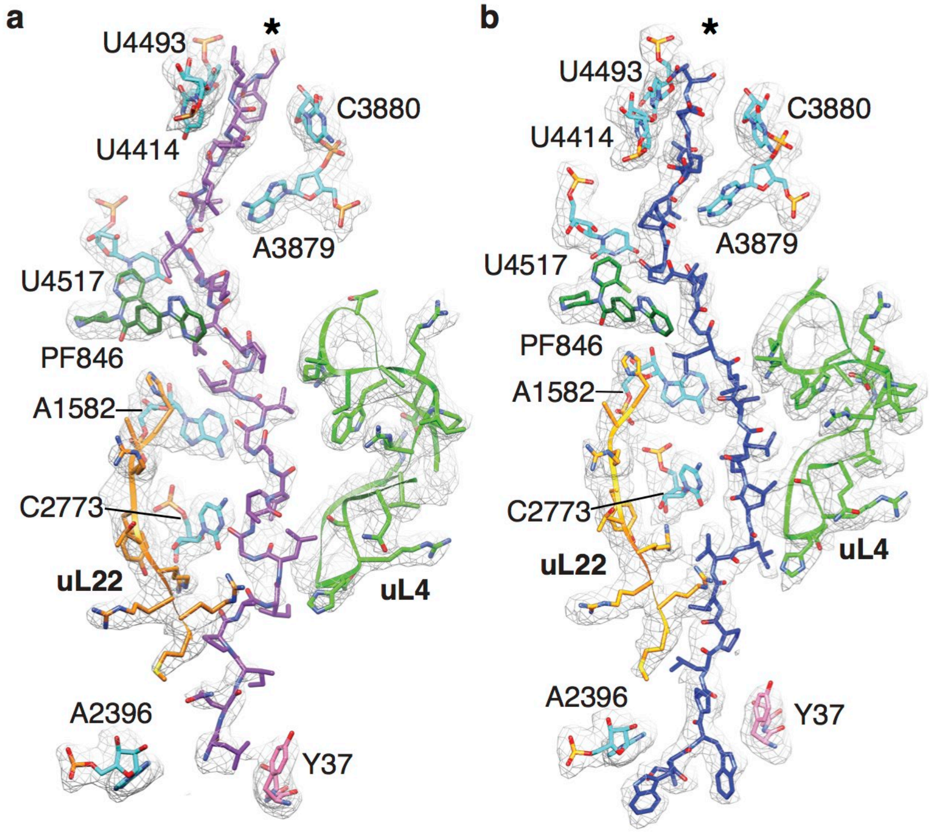

**Extended Data Fig. 10 | Interactions between the nascent chains and the ribosome exit tunnel. a-b,** Density map in the region where the (**a**) CDH1 (purple) and (**b**) PCSK9 (blue) nascent chains interact with the ribosome exit tunnel. Map density is shown in mesh, ribosomal proteins and rRNA nucleotides that have interaction with NC are shown in stick form (uL4 in green, uL22 in orange, eL39 in pink and 28S rRNA in cyan). The atomic models of the nascent chains are also shown in stick representation.

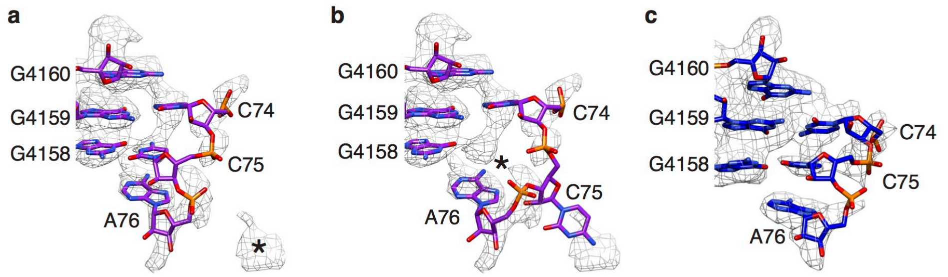

**Extended Data Fig. 11 | Poor base pairing of the 3’-CCA end of A/P NC-tRNA in PCSK9-stalled and USO1-stalled RNCs. a**, Interactions in the PCSK9-RNC between the 3’-CCA end of A/P tRNA and 28S rRNA nucleotides in the P-loop, with high-resolution structure of 3’-CCA end of peptidyl-tRNA (PDB: 1vy4) docked as a rigid body. **b**, A/P tRNA nucleotide C75 was flipped to nearby density without pairing with the P-loop (starting model from PDB: 1vy4). **c**, Interactions in the USO1-RNC, as in panel (**a**).

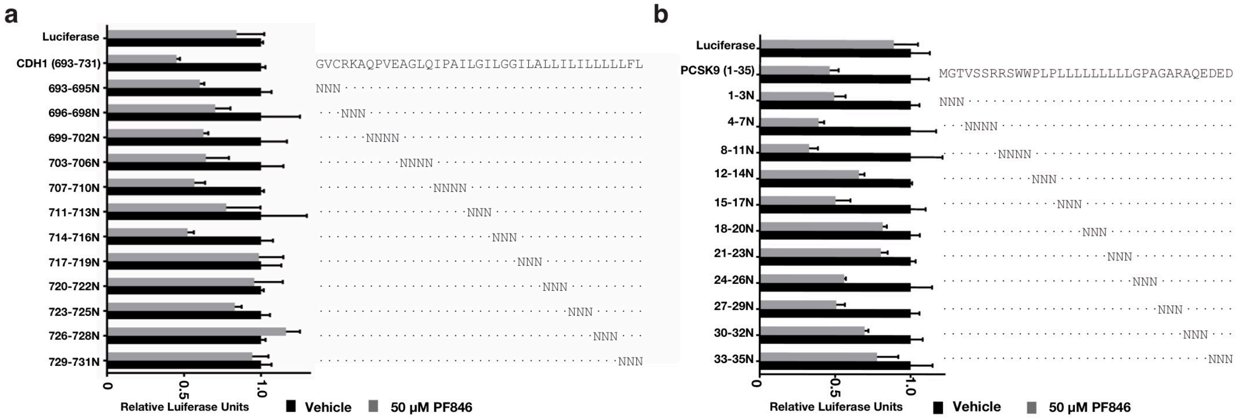

**Extended Data Fig. 12 | Asparagine scans of the CDH1 and PCSK9 NC sequences for effects on PF846-induced stalling. a**, Luciferase reporter assays using CDH1-derived NC sequences. CDH1 sequence is shown in full at the top, and locations of Asparagine (N) insertions are indicated. Luciferase, Luciferase-only control sequence. Reactions were carried out in the absence (black bar) or presence (grey bar) of PF846, in triplicate with standard deviations shown. **b**, Luciferase reporter assays using PCSK9-derived NC sequences. PCSK9 sequence is shown in full at the top, and locations of Asparagine (N) insertions are indicated. Luciferase, Luciferase-only control sequence. Reactions were carried out in the absence (black bar) or presence (grey bar) of PF846, in triplicate with standard deviations shown.

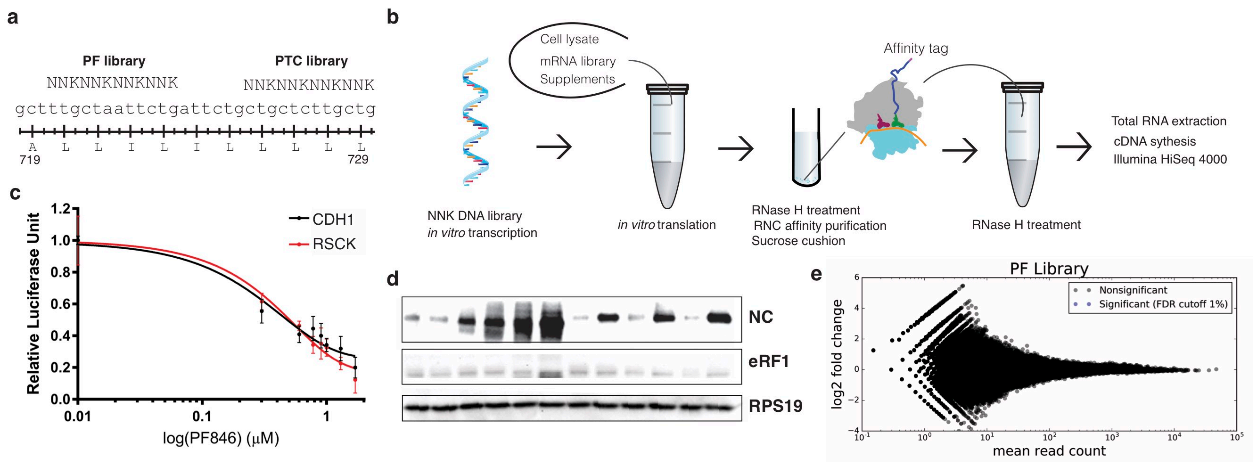

**Extended Data Fig. 13 | Selection of stalling sequences from mRNA libraries. a**, Schematic of mRNA library designs, with randomized locations in the CDH1 stalling sequence indicated. Numbers indicate amino acids in the CDH1 sequence. **b**, Protocol for mRNA library selections using *in vitro* translation reactions. RNase H treatment was carried out first during RNC binding and purification on anti-FLAG beads, and a second time after RNC release from the beads. **c**, Luciferase activity of translation reactions with luciferase reporters containing the CDH1 stalling sequence with the WT or RSCK motif near the PTC, as a function of PF846 concentration. Reactions were carried out in triplicate, with standard deviation shown. **d**, Western blot of affinity-purified stalled CDH1-derived nascent chains. Reactions in the absence or presence of 50 μM PF846 are shown. A luciferase reporter with no PF846-dependent stalling sequence but with an N-terminal 3x-FLAG tag is shown as a control (lanes 1, 2). Lanes 3 and 4 are WT CDH1 sequence; lanes 5 and 6 are CDH1-RSCK; lanes 7 and 8 are CDH1-CVT*; lanes 9 and 10 are CDH1-DPC*, and lanes 11 and 12 are CDH1-NVI*. Western blot of eRF1 for each sample is also shown, along with a Western of RPS19 as a loading control. **e**, MA plot of sequences enriched for PF846-induced translation elongation stalling, compared to translation reactions in the absence of PF846. Sequences were enriched from the mRNA library of CDH1-derived nascent chains with four amino acid positions randomized near the predicted location of PF846 binding in the ribosome exit tunnel. $Log_2$-fold enrichment of sequences is plotted against total read count, for experiments carried out in duplicate. No sequences were enriched with an adjusted *P*-value < 0.01.

**Extended Data Table 1 | Cryo-EM data collection, refinement and validation statistics of CDH1-RNC**

| | CDH1-RNC_AP tRNA | CDH1-RNC_AA tRNA | CDH1-RNC_PP tRNA |
|---|---|---|---|
| **Data collection and processing** | | | |
| Voltage (kV) | 300 | 300 | 300 |
| Electron exposure (e–/Å$^2$) | 50 | 50 | 50 |
| Defocus range (μm) | 1.0 to 2.0 | 1.0 to 2.0 | 1.0 to 2.0 |
| Pixel size (Å) | 1.15 | 1.15 | 1.15 |
| Symmetry imposed | C1 | C1 | C1 |
| Final particle images (no.) | 73,110 | 10,350 | 16,594 |
| Map resolution (Å) | 3.0 Å | 3.9 Å | 3.4 Å |
| FSC threshold | 0.143 | 0.143 | 0.143 |
| **Refinement** | | | |
| Initial model used (PDB code) | | | |
| 40S subunit | 6ek0 | 6ek0 | 5aj0 |
| 60S subunit | 5aj0 | 5aj0 | 5aj0 |
| tRNA | 3j7r | 3j7r | 5aj0 |
| Map sharpening *B* factor (Å$^2$) | -71 | -73 | -62 |
| R.m.s. deviations | | | |
| Bond lengths (Å) | 0.02 | 0.019 | 0.008 |
| Bond angles (°) | 1.199 | 1.188 | 1.081 |
| Validation | | | |
| Clashscore | 5.98 | 7.41 | 5.84 |
| Ramachandran plot | | | |
| Favored (%) | 88.87 | 87.9 | 88.81 |
| Allowed (%) | 10.63 | 11.76 | 10.71 |
| Outliers (%) | 0.5 | 0.34 | 0.48 |

**Extended Data Table 2 | Cryo-EM data collection, refinement and validation statistics of PCSK9-RNC**

| | PCSK9-RNC_AP tRNA | PCSK9-RNC_AA tRNA | PCSK9-RNC_PP tRNA |
|---|---|---|---|
| **Data collection and processing** | | | |
| Voltage (kV) | 300 | 300 | 300 |
| Electron exposure (e–/Å$^2$) | 50 | 50 | 50 |
| Defocus range (μm) | 1.0 to 2.5 | 1.0 to 2.5 | 1.0 to 2.5 |
| Pixel size (Å) | 1.15 | 1.15 | 1.15 |
| Symmetry imposed | C1 | C1 | C1 |
| Final particle images (no.) | 43,666 | 9,564 | 7,214 |
| Map resolution (Å) | 3.08 Å | 3.72 Å | 3.9 Å |
| FSC threshold | 0.143 | 0.143 | 0.143 |
| **Refinement** | | | |
| Initial model used (PDB code) | | | |
| 40S subunit | 6ek0 | 6ek0 | 5aj0 |
| 60S subunit | 5aj0 | 5aj0 | 5aj0 |
| tRNA | 3j7r | 3j7r | 5aj0 |
| Map sharpening *B* factor (Å$^2$) | -75 | -77 | -80 |
| R.m.s. deviations | | | |
| Bond lengths (Å) | 0.019 | 0.020 | 0.009 |
| Bond angles (°) | 1.190 | 1.296 | 1.222 |
| Validation | | | |
| Clashscore | 5.92 | 7.36 | 7.33 |
| Ramachandran plot | | | |
| Favored (%) | 89.56 | 87.47 | 87.23 |
| Allowed (%) | 10.03 | 12.21 | 12.20 |
| Outliers (%) | 0.41 | 0.32 | 0.58 |

**Extended Data Table 3 | Cryo-EM data collection statistics of PCSK9s-RNC**

| | PCSK9s-RNC_AP tRNA | PCSK9s-RNC_PP tRNA |
|---|---|---|
| **Data collection and processing** | | |
| Voltage (kV) | 300 | 300 |
| Electron exposure (e–/Å$^2$) | 50 | 50 |
| Defocus range (μm) | 1.0 to 2.5 | 1.0 to 2.5 |
| Pixel size (Å) | 1.22 | 1.22 |
| Symmetry imposed | C1 | C1 |
| Final particle images (no.) | 8,631 | 8,433 |
| Map resolution (Å) | 4.68 Å | 4.57 Å |
| FSC threshold | 0.143 | 0.143 |

**Extended Data Table 4 | Cryo-EM data collection, refinement and validation statistics of USO1-RNC**

| | USO1-RNC |
|---|---|
| **Data collection and processing** | |
| Voltage (kV) | 300 |
| Electron exposure (e–/Å$^2$) | 55 |
| Defocus range (μm) | 1.5 to 2.5 |
| Pixel size (Å) | 1.15 |
| Symmetry imposed | C1 |
| Final particle images (no.) | 38,314 |
| Map resolution (Å) | 3.5 Å |
| FSC threshold | 0.143 |
| **Refinement** | |
| Initial model used (PDB code) | |
| 40S subunit | 6ek0 |
| 60S subunit | 5aj0 |
| tRNA | 3j7r |
| Map sharpening *B* factor (Å$^2$) | -91 |
| R.m.s. deviations | |
| Bond lengths (Å) | 0.018 |
| Bond angles (°) | 1.205 |
| Validation | |
| Clashscore | 6.84 |
| Ramachandran plot | |
| Favored (%) | 89.33 |
| Allowed (%) | 10.34 |
| Outliers (%) | 0.33 |

### Extended Data Table 5 | Calculated energies of various conformations of PF846.

| Molecule name | Energy (kcal/mol) |
| --- | --- |
| Optimized_X-ray_molecule_1 | -1110331.97 |
| Optimized_X-ray_molecule_2 | -1110333.04 |
| Pose 1 | -1110331.49 |
| Pose 2 | -1110331.64 |
| Pose 3 | -1110333.24 |
| Pose 4 | -1110333.59 |
